# Identification of PKN2 and MOB4 as Coordinators of Collective Cell Migration

**DOI:** 10.1101/2025.02.14.638398

**Authors:** Artem I. Fokin, Yueying Lin, Dmitry Y. Guschin, Hsiang-Ying Chen, John James, Jie Yan, Pascal Silberzan, Alexis M. Gautreau

**Author notes:** Scientific Center for Translational Medicine, Sirius University of Science and Technology. 254349 Sirius, Russian Federation (Dmitry Y. Guschin).

## Abstract

In animals, collective cell migration is critical during development and adult life for repairing organs. It remains, however, poorly understood compared with single cell migration. The polymerization of branched actin by the RAC1-WAVE-Arp2/3 pathway is established to power membrane protrusions at the front of migrating cells, but also to maintain cell junctions in epithelial monolayers. Here we have identified novel regulators of collective cell migration using a two-pronged approach: candidates were extracted from publicly available RAC1-WAVE-Arp2/3 dependency maps and screened in a second step using CRISPR/Cas9 genetic inactivation. In a wound healing assay, PKN2 knockout (KO) MCF10A cells display decreased collective migration due to destabilization of adherens junctions, whereas MOB4 KO cells display increased collective migration with a loss of migration orientation. Upon wound healing, PKN2 relocalizes to lateral junctions and maintains coordinated migration in the monolayer, whereas MOB4 relocalizes to the front edge of leader and follower cells collectively migrating towards the wound. The role of MOB4 in controlling collective migration requires YAP1, since MOB4 KO cells fail to activate YAP1 and their phenotype is rescued by constitutively active YAP1. Together, these findings reveal two complementary activities required for coordinating cells in collective migration.

## 1. INTRODUCTION

Collective cell migration is essential to the development and adult life of animals, starting from gastrulation, continuing later on during organogenesis and healing of injuries throughout life ^[1]^. It is also important for pathological processes, such as cancer progression and tumor cell invasion ^[2]^. Collective cell migration displays similarities with single cell migration. For example, cells at the front of a migrating epithelium develop at their leading edge an adherent membrane protrusion called a lamellipodium, like single migrating cells ^[3,4]^. The lamellipodium is not restricted to leader cells. It is also present in follower cells, even if they are surrounded by cell-cell junctions. The lamellipodium in this case is referred to as cryptic, because it is hidden beneath the cell located immediately ahead ^[5–8]^.

Lamellipodia are the results of the pushing force exerted by the polymerization of branched actin through the Arp2/3 complex ^[9,10]^. Cortical branched actin also pushes against the cell-cell junctions of epithelial cells and this activity has been demonstrated to be critical for the establishment and maintenance of adherens junctions ^[11–13]^. Cell-cell junctions experience opposing forces: pulling forces due to myosin-mediated contractility and pushing forces due to Arp2/3-mediated branched actin. Since excessive contractility can induce bulges between adjacent cells, the Arp2/3-mediated pushing forces therefore helps to repair junctions by affixing neighboring cells ^[14,15]^. The same cortical branched actin pathway that depends on the small GTPase RAC1 and the WAVE complex is therefore simultaneously required for lamellipodial protrusions and cell-cell junctions in the collective migration of epithelial cells ^[16–19]^. This implies that the same pathway generating branched actin polymerization is differentially regulated at these two locations to give rise to two different structures. But how this differential regulation is achieved is currently not known.

Epithelial cells stop migrating and proliferating when they reach a high cell density, in a process known as jamming ^[20–22]^. The arrested monolayer resumes migration and proliferation upon unjamming, which can be mechanically induced either in an isotropic manner using biaxial stretching or a hypo-osmotic shock, or in an anisotropic manner by wounding the monolayer ^[23– 26]^. The induced migration of cells is then coordinated over large distances, a process referred to as fluidization of the epithelial monolayer ^[27,28]^ . In wound healing, collective cell migration is directed towards the wound. Leader cells facing the wound instruct follower cells to migrate in the same direction through mechanotransduction across cell-cell junctions ^[29]^. The mechanotransducer vinculin, well established to connect actin filaments to integrin-mediated focal adhesions, also connects actin filaments to E-cadherin-mediated adherens junctions ^[30]^. The NF2 tumor suppressor, which is associated with cell-cell junctions in a jammed epithelium and relocalizes to the cytosol upon unjamming, appears to transmit wound sensing into activation of the GTPase RAC1 and the YAP1 transcription factor ^[31,32]^. In wound healing, the mechanosensitive YAP1 is activated in leader cells and induces the expression of genes associated with proliferation and adhesion ^[33–35]^.

To gain insights into the mechanisms of cell coordination in collective migration, we interrogated publicly available dependency maps using components of the RAC1-WAVE-Arp2/3 pathway, since this pathway is involved in establishing and maintaining lamellipodial protrusions and cell junctions, and performed a miniscreen using CRISPR/Cas9 inactivation of candidate genes. This two-pronged approach allowed us to identify two previously unknown regulators of cell migration: PKN2 that maintains epithelial cohesiveness in collectively migrating cells, and MOB4 that restrains collective migration and that indicates the proper direction of migration towards the wound.

## 2. RESULTS

### 2.1 Selection of Candidate Genes and Isolation of Knockout Clones

The RAC1-WAVE-Arp2/3 pathway is well established to control polymerization of branched actin and cell migration. The same pathway, however, also controls progression into the cell cycle and hence cell proliferation ^[36–38]^. To identify novel effectors of the RAC1-WAVE-Arp2/3 pathway, we used the latter property and exploited the public database depmap.org, reporting dependency maps ^[39]^. The depmap consortium performed, in many cell lines, the following genetic screen to identify genes that are critical for growth: a library of gRNAs is transduced into a cell line and analyzed by Next Generation Sequencing (NGS), first at the time of transduction and then after 16 doubling of the cell population (Figure 1A). The comparison of the two NGS allows identifying gRNAs that are significantly depleted, corresponding to essential genes, or enriched, corresponding to tumor suppressor genes. Given that this genetic screen has already been performed on more than 500 cell lines, genes encoding different signaling intermediates in the same pathway showed statistical correlation, i.e. they were essential in some cell lines and non-essential in others. We retrieved the genes correlated with RAC1 and with the subunits of WAVE and Arp2/3 complexes, from depmap.org.

**Figure 1.**
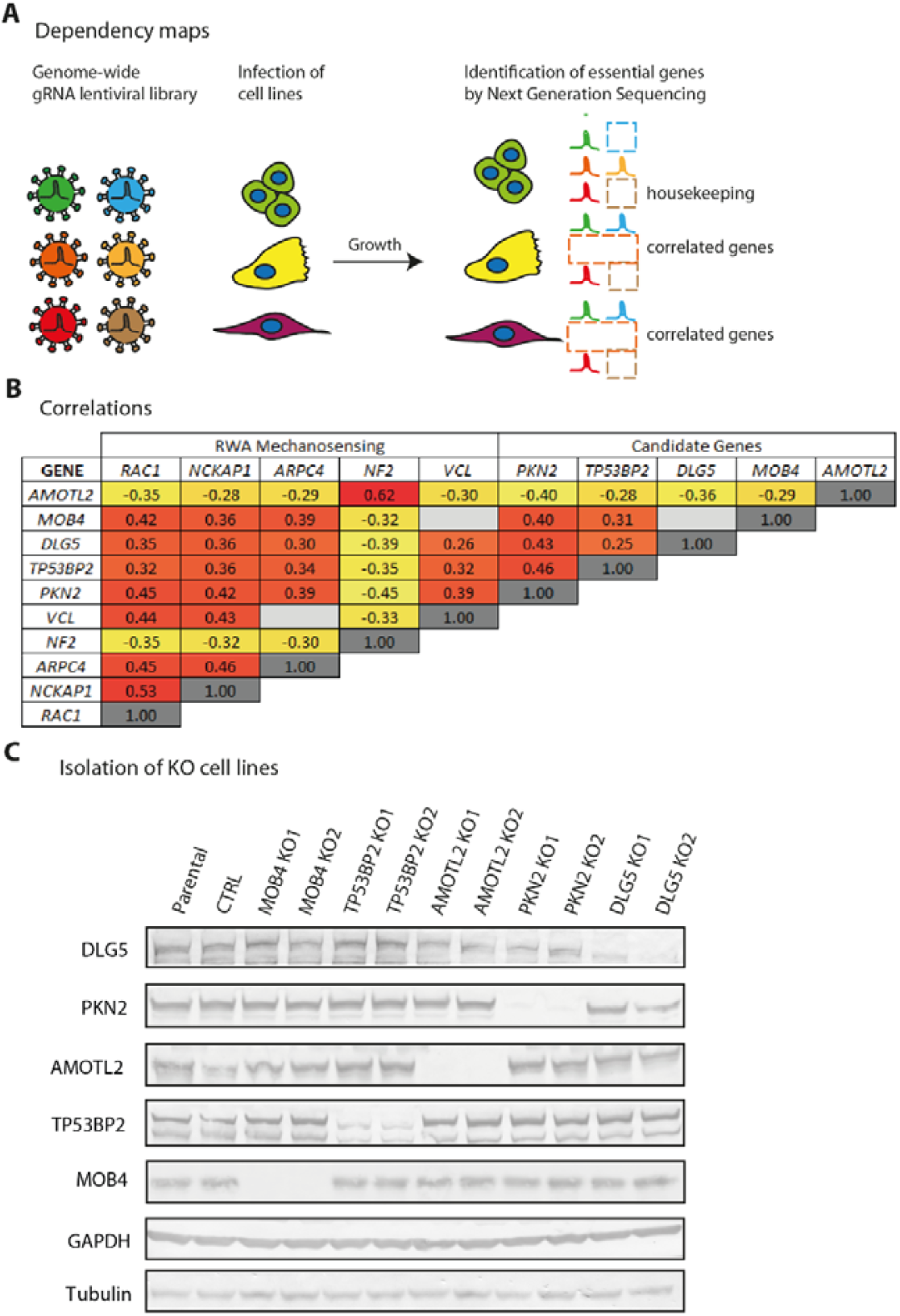
Selection of candidate genes and isolation of Knock-Out (KO) clones. A) Principle of dependency maps. Publicly available results reveal essential genes in more than 500 cell lines. Genes that are required in the same subset of lines are correlated and may function in the same pathway. B) Matrix reporting gene correlations between known genes of the mechanosensing RAC1-WAVE-Arp2/3 pathway (RWA) and five new candidate genes associated with cell-cell junctions. Hot colors highlight high correlations. *RAC1* encodes the small GTPase RAC1; *NCKAP1* and *ARPC4* encode subunits of the WAVE and Arp2/3 complex, respectively. *NF2* encodes the tumor suppressor protein Merlin, and *VCL* encodes the mechanotransducer protein vinculin. C) KO of the five candidate genes was obtained in the MCF10A breast epithelial cell line. Two independent clones of each were analyzed by Western blots using the indicated antibodies.

As expected, the genes of the RAC1-WAVE-Arp2/3 pathway showed a positive correlation (Figure 1B). The gene encoding vinculin, a mechanotransducer associated with Arp2/3 ^[40,41]^, showed a similar correlation. In contrast, *NF2*, a tumor suppressor gene known to antagonize the RAC1 pathway ^[42,43]^, was anti-correlated with all of the above factors. These correlations between known effectors of the RAC1-WAVE-Arp2/3 pathway confirmed that effectors of this molecular pathway can be deduced from dependency maps. Most of the genes correlated with the RAC1-WAVE-Arp2/3 pathway that were extracted from dependency maps belonged to the integrin pathway. These were effectors linking the specific interaction of integrins with the extracellular matrix to the polymerization of branched actin ^[44]^. For example, among the correlated genes, we found Focal Adhesion Kinase and Kindlin-2, two proteins in the integrin pathway that interact with the Arp2/3 complex ^[45,46]^. However, the genes of the integrin pathway were not our main interest in this study and were therefore not investigated further.

Given our interest in collective cell migration, we rather focused on genes encoding proteins that were physically or functionally associated with cell-cell junctions. This allowed us to select five genes: *PKN2, TP53BP2, DLG5, MOB4* and *AMOTL2*. Five genes is a tractable number of candidate genes for subsequent CRISPR/Cas9 mediated gene inactivation. The *PKN2* gene encodes the serine/threonine protein kinase N2, PKN2, also known as PRK2. PKN2 is an effector of both RHOA and RAC1 GTPases that regulates the formation of cell junctions ^[47,48]^. PKN2 was found in the E-cadherin interactome ^[49]^. The *TP53BP2* gene encodes a p53 binding protein, TP53BP2, which is also known as ASPP2. TP53BP2 binds to E-cadherin and β-catenin and regulates epithelial to mesenchymal transition (EMT) ^[50]^. TP53BP2 also associates with the NF2 tumor suppressor ^[51]^. TP53BP2 maintains the integrity of stressed epithelia and inhibits cell migration and proliferation ^[52,53]^. *DLG5* encodes the adapter protein Disks large homologue 5, a conserved member of the MAGUK family. DLG5 interacts with β-catenin ^[54]^ and maintains the integrity of epithelial monolayers ^[55]^. DLG5 depletion induces EMT and increases cell migration and proliferation ^[56,57]^. *MOB4* encodes a protein of the phocein family. MOB4 regulates the mechanosensitive YAP1 pathway by associating with the MST4 kinase ^[58]^ and contributes to a large assembly containing the PP2A phosphatase called the STRIPAK complex ^[59,60]^. The negatively correlated gene *AMOTL2*, encodes an Angiomotin family protein, which can be an oncogene or a tumor suppressor depending on the cancer type ^[61,62]^. AMOTL2 localizes to cell-cell junctions and regulates the NF2 tumor suppressive activity ^[63]^.

For fast and efficient knock-out (KO) of these candidate genes, we transfected the cells with plasmids encoding Cas9 and three gRNAs: two that introduce double-strand breaks in the target gene, thus inducing a deletion, and one that targets the gene encoding the essential Na+/K+ antiporter ATP1A1 at the position of its ouabain binding site. Ouabain thus selects cells carrying mutated *ATP1A1*, where the single double-strand break has been repaired by non-homologous end joining, in such a way that the antiporter is functional, but insensitive to ouabain ^[64]^. This efficient selection renders deletion screening fast (with 10 to 90 % success rate) and permitted us to retrieve KO clones from the MCF10A breast epithelial cell line for each five target genes. We selected two KO clones for each gene, whose alleles were sequenced to identify precise deletions, resulting most often in a frameshift (Figure S1). All KO clones were further validated by Western blots (Figure 1C). We compared these KO clones with parental MCF10A cells and a pool of ouabain-resistant cells referred to below as control cells.

### 2.2 Screening Candidate Genes for their Role in Collective Cell Migration

We then tested these KO cell lines in a model of wound healing, obtained by lifting inserts to create a cell-free area^[65,66]^. MCF10A cells migrate and close the wound with a smooth progressing edge. To quantitatively analyze the coordination of collectively migrating cells, we applied particle image velocimetry (PIV) to movies acquired with phase contrast ^[67]^. PIV consists in dynamic mapping the velocity field over the entire field of view, via local correlations between successive images. The field of displacement vectors varied over time and allowed extracting two major parameters: the speed of cells and the order parameter that reflects the migration direction towards the wound ^[68]^. These parameters are then represented as heat maps to capture their evolution in time and space (Figure 2A). We also calculated velocity correlation length, as a measure of the coordination of cell movement, which is independent from the migration direction.

**Figure 2.**
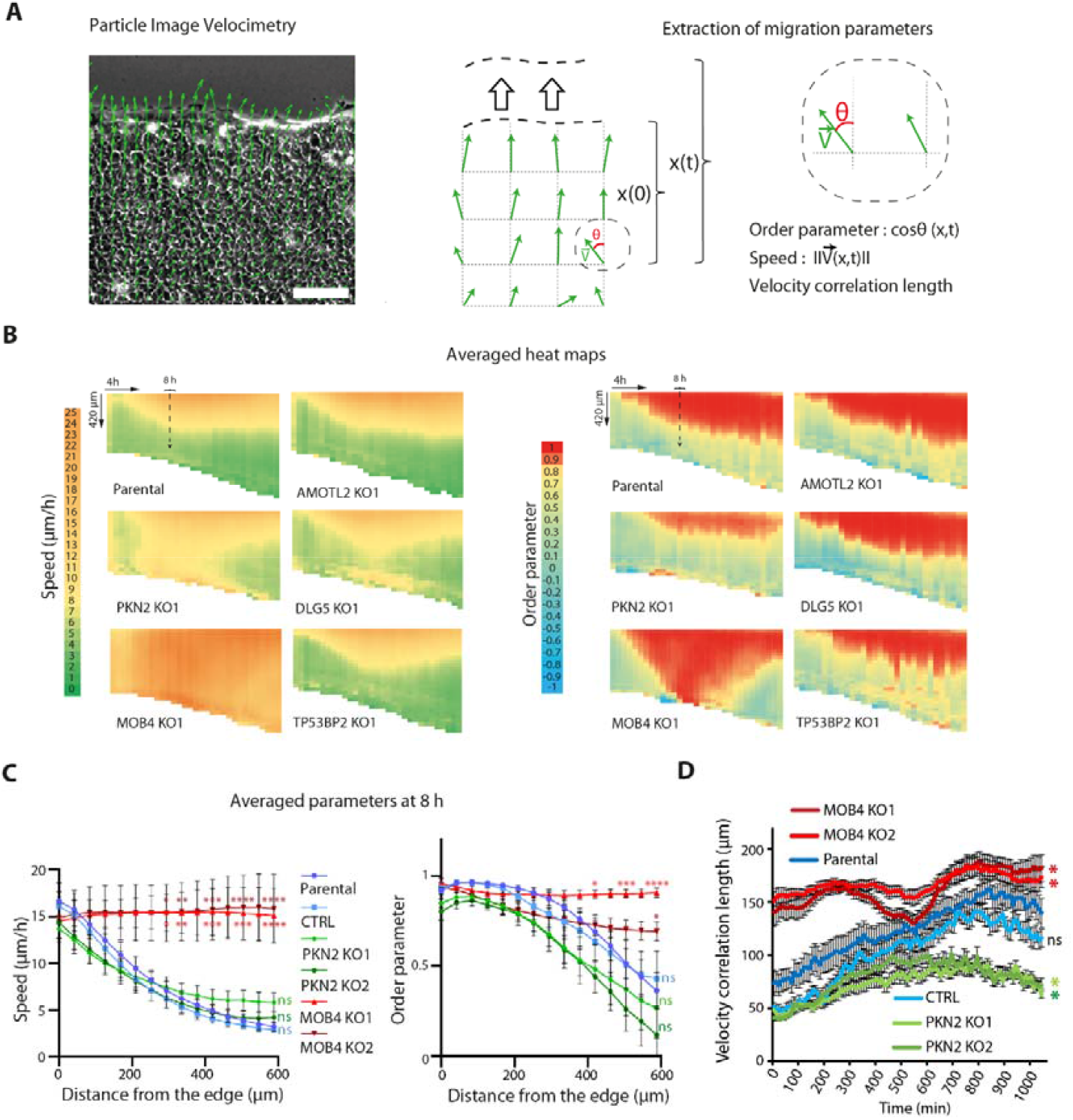
Identification of *PKN2* and *MOB4* as genes that control the coordination of collective cell migration. A) Fields of displacement vectors are obtained by Particle Image Velocimetry (PIV) of the wound healing movie. The vector field is superimposed to the phase contrast image of the MCF10A monolayer. Definition of migration parameters extracted from vector fields. Scale bar: 100 µm. B) Speed and order parameter are plotted as heat maps across time and space. This heat map representation shows the monolayer front facing the wound on top and time 0 on the left. For one KO clone of each candidate gene, averaged heat maps of eight movies from two independent experiments are plotted. Each independent experiment gave similar results. The dashed line represents the time 8 h. C) Local speed and order are averaged at 8 h after wounding from the 12 fields of view of the three biological repeats. Mean ± SEM, two-way ANOVA. D) Velocity correlation length calculated from 11 fields of view from the three biological repeats is plotted as a function of time. Areas under curve are compared by one-way ANOVA, mean ± SEM. * p<0.05, ** p<0.01, *** p<0.001, **** p<0.0001, ns non-significant.

Parental cells or ouabain-resistant control cells exhibited a stereotypical response to the wound (Figure 2B, Figure S2): leader cells began migrating towards the wound; the extent of follower cells collectively migrating towards the wound expanded over time, up to about 500 μm in depth. This gradient of collectively migrating cells can be read from heat maps of speed or order parameter. The genetic inactivation of three selected genes, *DLG5, AMOTL2* and *TP53BP2*, did not alter heat maps of speed or order parameter. In contrast, two of our selected genes, PKN2 and MOB4, affected collective migration, when being knocked out (Figure 2B, Figure S2 for the second set of clones).

Migration of PKN2 KO cells does not appear as coordinated as in parental cells (Video S1). Overall, PKN2 KO marginally decreased both speed and order parameter. At the arbitrary time point of 8 h after wounding, chosen for comparison throughout the text, differences of speed and order parameters were just a trend that was not statistically significant (Figure 2C, Figure S2). Unlike control cells, however, the velocity correlation length of PKN2 KO cells failed to increase over time and this difference was statistically significant (Figure 2D, Figure S2). We confirmed these results by manual tracking of PKN2 KO cells at different distances from the wound, using the same recordings (Video S2). Individual trajectories of PKN2 KO cells did not exhibit the steep gradient response of parental cells. Speed and distance traveled was more homogenous in PKN2 KO cells between front and back than in parental cells (Figure S2). Directional persistence towards the wound was also decreased compared with parental cells throughout the migrating monolayer (Figure S2). The two methods, PIV and manual tracking, thus converge to suggest that the PKN2 protein promotes collective cell migration. PIV generates more data in less time and avoids the bias of having to pick the cells to track. PIV thus remains the method of choice to analyze collective cell migration.

Migration of MOB4 KO cells in wound healing appears more collective than parental cells: the territory of collectively migrating KO cells is not restricted to the first rows of cells facing the wound as in parental cells, but the migration direction of MOB4 KO cells appear erratic (Video S3, Video S4 for further examples). A gradient of speed was simply not observed upon MOB4 inactivation: MOB4 KO cells exhibited high speed throughout the field of view (Figure 2B, Figure S3). MOB4 KO cells indeed displayed increased collective migration, as evidenced by the increased velocity correlation length compared with that of controls (Figure 2D). Erratic orientation of collective migration translated into irregular variations of the order parameter among individual fields of view (Figure S3), an effect that was alleviated upon averaging the different fields of view (Figure 2B). The complex phenotype of MOB4 KO cells suggests that the MOB4 protein might both restrict collective cell migration and indicate the direction that collectively migrating cells should take.

Our miniscreen thus revealed a positive and a negative regulator of collective migration, PKN2 and MOB4, respectively. Yet genetic inactivation of these two opposite regulators resulted in a slightly decreased efficiency of the monolayer at closing the wound in both cases (Figure S4). For the sake of completeness, we then sought to characterize the effect of these two genes on single cell migration. PKN2 KO cells were overall not significantly different from controls (Figure S5, Video S5). In contrast, MOB4 KO cells exhibited increased directional persistence, decreased speed and Mean Square Displacement (MSD) compared with parental cells in the single cell assay (Figure S5, Video S6).

### 2.3 PKN2 and MOB4 Associate With Front or Lateral Junctions

To better understand the roles of PKN2 and MOB4 in collective cell migration, we generated stable MCF10A cell lines expressing either GFP-PKN2 or GFP-MOB4. The two proteins appeared mostly cytosolic. Upon wounding, a small fraction of both proteins translocates to junctions of migrating cells (Video S7). The membrane pools of PKN2 and MOB4 colocalize with WAVE2 (Figure S6). Moreover, when PKN2 and MOB4 were immunoprecipitated, a small amount of WAVE complex, as shown by the WAVE2, NCKAP1 and ABI1 subunits, was coprecipitated (Figure S6).

All junctions were not equally stained by GFP-PKN2 and GFP-MOB4. A closer examination of the cell junctions stained by GFP-PKN2 and GFP-MOB4 revealed that most junctions decorated with PKN2 were orthogonal to the wound edge, whereas most junctions decorated with MOB4 were parallel to the edge (Figure 3A). The enrichment of GFP-MOB4 at junctions parallel to the edge might occur either at the front or the back of cells. To distinguish between these two possibilities, we mixed GFP-MOB4 expressing cells to non-fluorescent parental MCF10A cells and observed that the front of fluorescent cells was stained (Video S8), suggesting that MOB4 is present in cryptic lamellipodia. We then sought to confirm these observations using immunofluorescence of endogenous PKN2 and MOB4. Using validated antibodies that do not stain the corresponding KO cell lines (Figure S7), we indeed observed that endogenous PKN2 decorated lateral cell junctions, whereas MOB4 decorated front cell junctions (Figure 3B), like the corresponding GFP fusion proteins. In leader cells, MOB4 also stains the free edge facing the wound, confirming that MOB4 is a lamellipodial protein.

**Figure 3.**
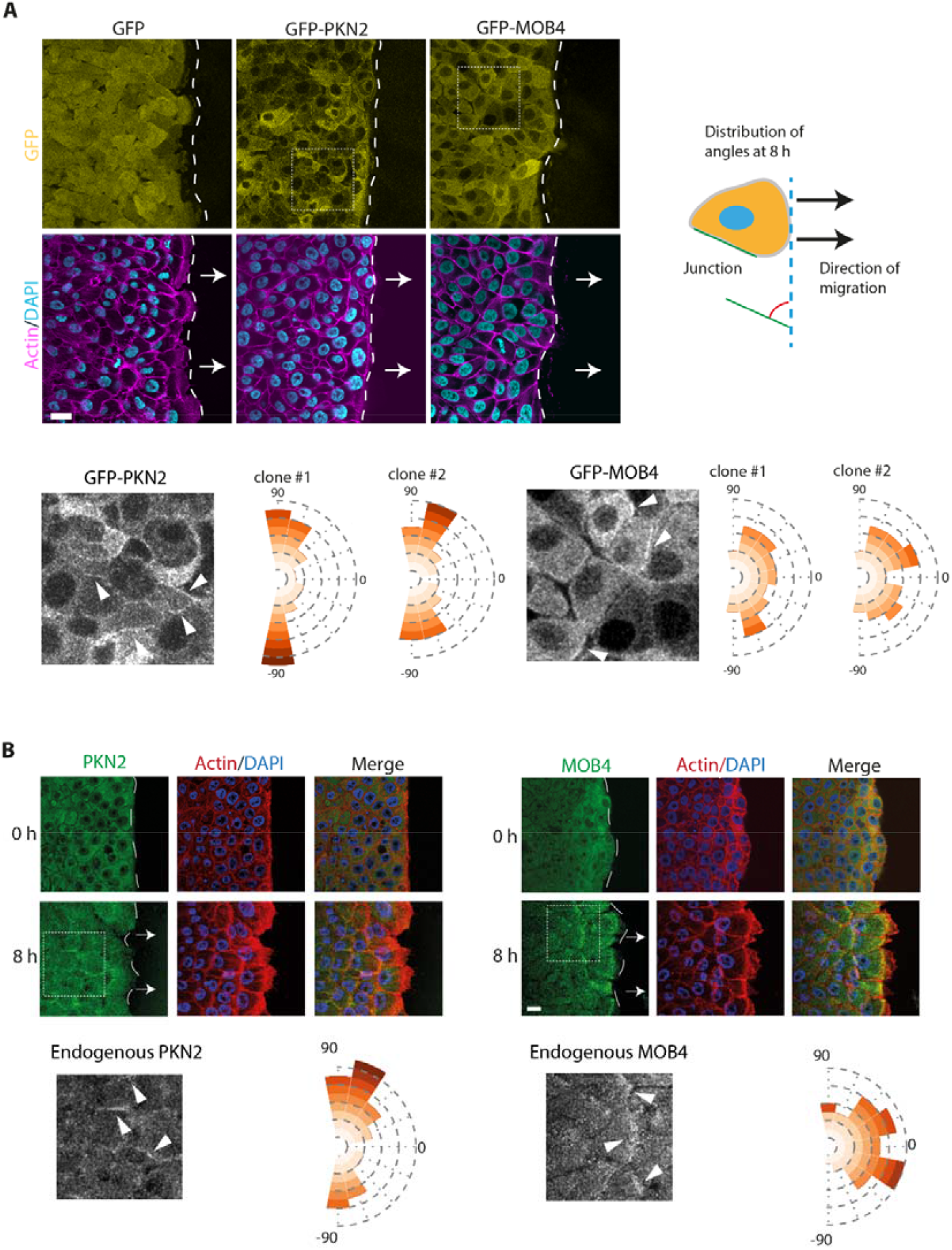
PKN2 and MOB4 differentially localize to the front and lateral edges of collectively migrating cells. A) Stable MCF10A lines expressing GFP-PKN2 or GFP-MOB4 are imaged by confocal microscopy. Filamentous actin is stained with phalloidin and nuclei with DAPI. Angle distribution of all cell junctions labeled by MOB4 or PKN2. In total, more than 100 junctions per cell line are assessed in three independent experiments. B) PKN2 and MOB4 immunofluorescence in MCF10A monolayers at time 0 or 8 h after wounding. Filamentous actin is stained with phalloidin and nuclei with DAPI. Angle distribution of all cell junctions labeled with MOB4 or PKN2. More than 50 junctions for each staining are assessed in two independent experiments. Confocal microscopy. Scale bars 20 µm.

### 2.4 PKN2 Controls Collective Migration by Maintaining Stable Cell Junctions

In an attempt to explain the decreased collective migration of PKN2 KO cells, we stained these cells for E-cadherin, the major cell-cell adhesion molecule of epithelial cells, which exerts calcium-dependent homotypic interactions between neighboring cells. PKN2 KO cells had a significantly decreased staining of E-cadherin at cell-cell junctions compared with parental cells (Figure 4A). This was not the case of MOB4 KO cells. When E-cadherin levels were examined by Western blot, no difference was observed between cell lines (Figure 4B), suggesting that E-cadherin is expressed, but not concentrated at cell junctions, in PKN2 KO cells. Because the RAC1-WAVE-Arp2/3 pathway generates branched actin at cell junctions, we also stained the three cell lines with antibodies targeting cortactin, which is a bona fide marker of Arp2/3-mediated branched actin ^[69]^ . Branched actin was not altered at cell junctions upon inactivation of PKN2 or MOB4 (Figure 4C,D). As a proxy to cell-cell adhesion, we used a biophysical assay consisting in measuring the rupture force, when an E-cadherin coated bead interacting with a cell is forced to detach from it ^[70]^. Consistent with the observed decreased pool of E-cadherin at the membrane of PKN2 KO cells, we observed decreased mechanical stability of E-cadherin coated beads interacting with PKN2 KO cells compared with parental cells (Figure 4E).

**Figure 4.**
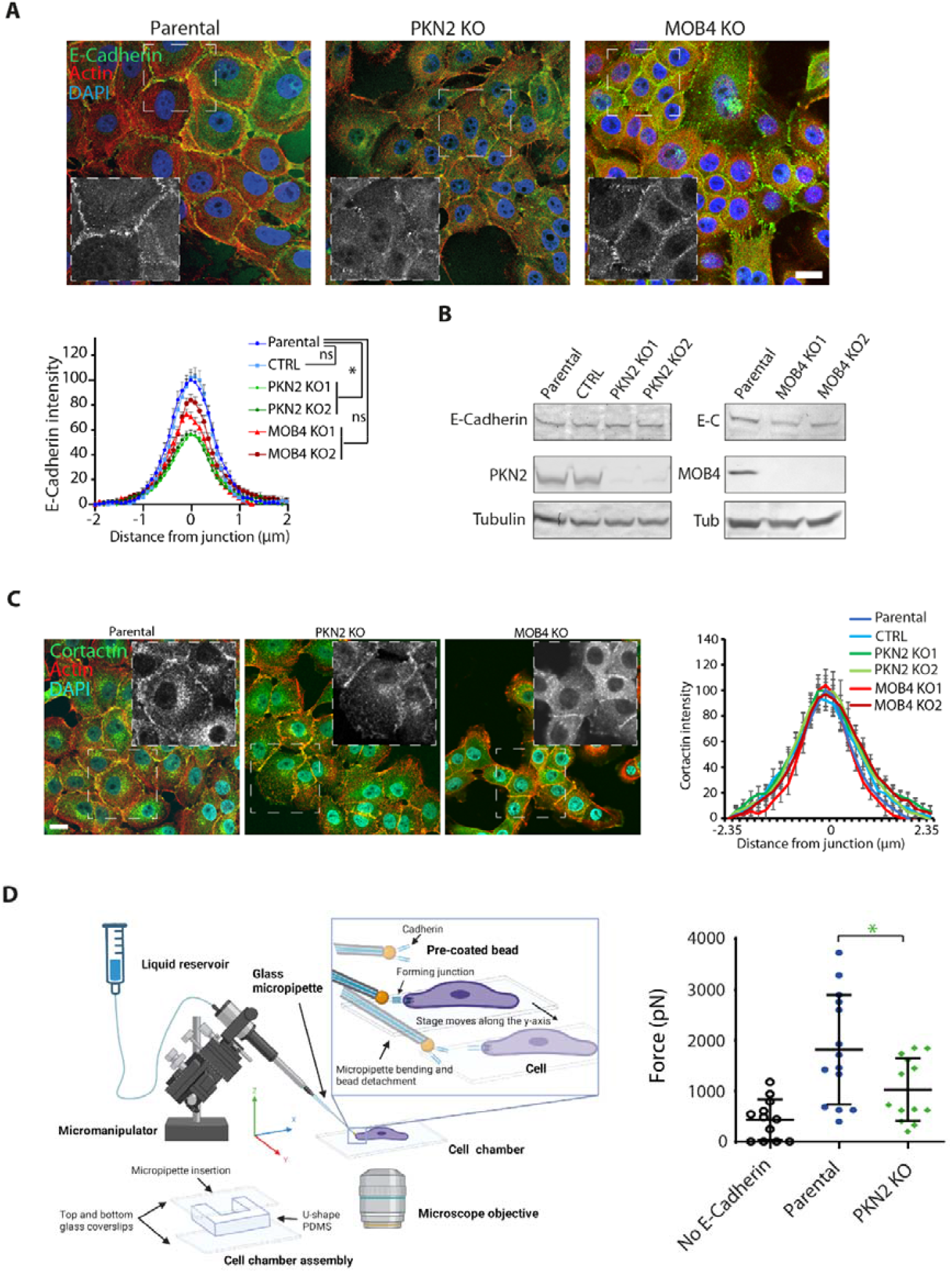
PKN2 KO cells display weakened adherens junctions. A) E-cadherin immunofluorescence of epithelial islets of PKN2, MOB4 KO and control cells. Filamentous actin is stained with phalloidin and nuclei with DAPI. Confocal microscopy. Scale bar 20 µm. Black-and-white inserts show E-cadherin staining in the magnified regions. Three independent experiments gave similar results. Intensity profiles taken perpendicularly to cell-cell junctions were averaged for a total of n=60 junctions. Mean ± SEM. Areas under the curve, one-way ANOVA. B) Western blots of E-cadherin, PKN2, MOB4 and tubulin as a loading control. C) Islets of PKN2 KO, MOB4 KO and parental cells are stained with antibodies to cortactin, phalloidin and DAPI. Confocal microscopy. Scale bar 20 µm. Black-and-white inserts show cortactin staining in the magnified regions. Intensity profiles of cortactin taken perpendicularly to cell-cell junctions were averaged for a total of n=20 junctions. Mean ± SEM. D) Measurements of the rupture force of E-cadherin coated beads in PKN2 KO and MCF10A parental cells. The control shown refers to force measurement when beads are not coated. The set-up used consists in a chamber composed of a U-shaped PDMS structure between two glass coverslips, where cells are seeded and manipulated. The liquid reservoir, when placed below the plane of the cells, allows aspirating beads. When the micromanipulator is used to displace the bead, the micropipette bends because of bead attachment to the cell until the bonds rupture. The force of the E-cadherin mediated interaction is calculated from the micropipette deflection, knowing the measured elasticity of the micropipette. Non-paired t-test. * p<0.05, ns non-significant.

In order to mimic the defective cell-cell adhesions of PKN2 KO cells, we treated parental cells with EDTA that chelates Ca^2+^ ions, or with EDTA and Ca^2+^ to neutralize the effect of EDTA. Upon wounding, we observed that EDTA-treated cells exhibited the same behavior as PKN2 KO cells: many cells at the front edge detached from the monolayer and migrated as single cells (Figure 5A, Video S9). Moreover, velocity correlation length in the monolayer was strongly reduced by EDTA treatment (Figure 5B). Heat maps of the order parameter showed a reduction compared with parental cells (Figure 5C, Figure S8). As expected, upon Ca^2+^ replenishment, all three effects seen upon EDTA addition were abolished. EDTA treatment thus phenocopies PKN2 inactivation, suggesting that the decreased coordination of PKN2 KO cells in collective migration might be due to impairment of adherens junctions formation.

**Figure 5.**
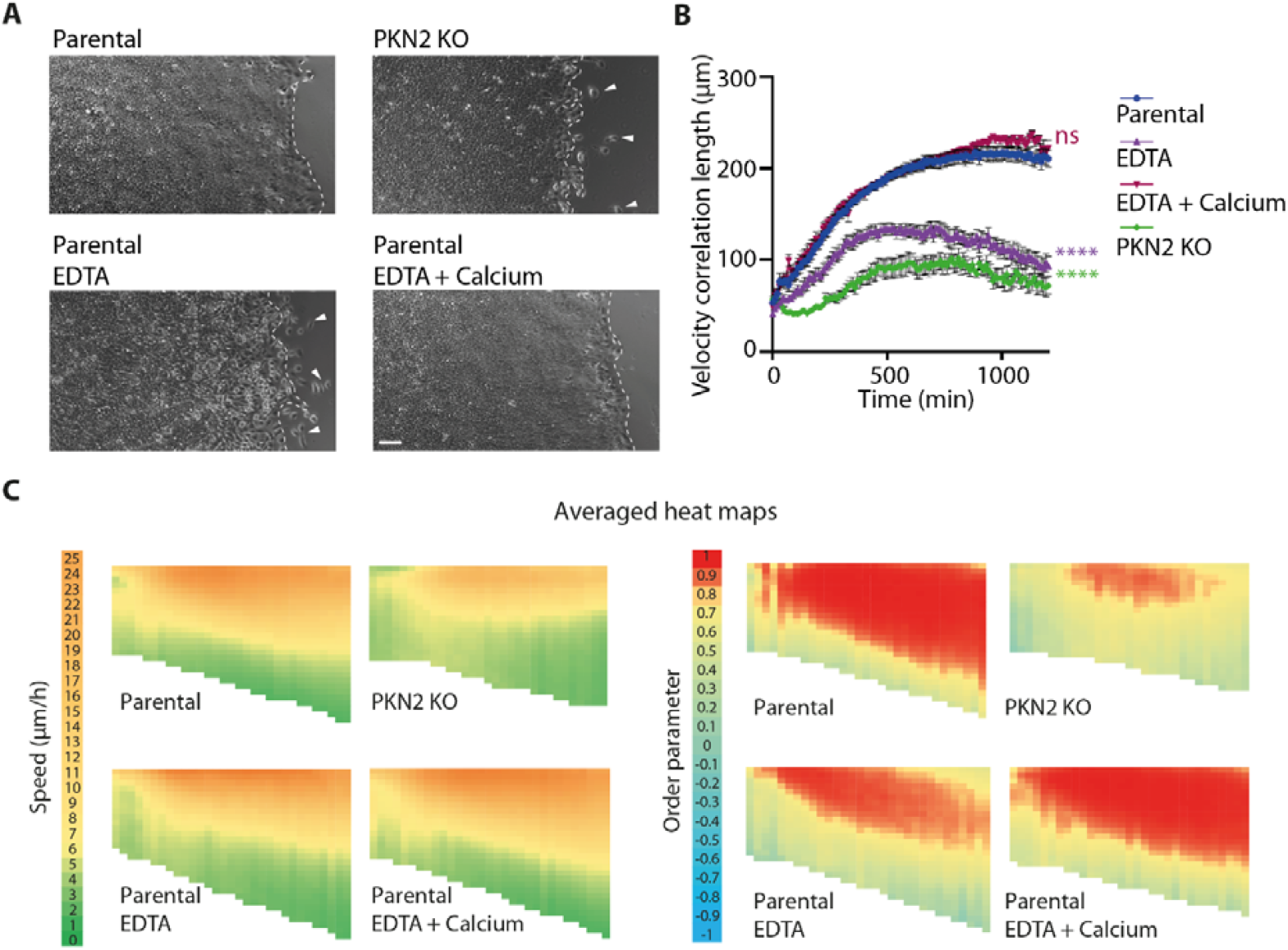
EDTA treatment, which weakens adherens junctions, phenocopies PKN2 KO cells. A) Wound healing of PKN2 KO cells or parental cells treated with 1 mM EDTA to destabilize cell-cell junctions or 1 mM EDTA with 1 mM CaCl_2_ to neutralize EDTA. EDTA-mediated destabilization of junctions phenocopies the PKN2 KO phenotype. Scale bar 100 µm. B) Velocity correlation length, calculated from 12 movies from two independent experiments, is plotted as a function of time. Areas under the curve are compared by one-way ANOVA, mean ± SEM, **** p<0.0001, ns non-significant. C) Heat maps of speed and order parameters are extracted from PIV analyses of 12 movies from two independent experiments per condition. Each independent experiment gave similar results.

### 2.5 MOB4 Controls Collective Migration by Activating YAP1

We then attempted to identify appropriate parameters to describe the complex phenotype of MOB4 KO cells. To represent increased collective migration and erratic cell polarity, we drew “streamlines” that concatenate successive displacement vectors obtained by PIV (Figure 6A). These streamlines were dramatically elongated in MOB4 KO cells compared with parental cells in line with the increased territory of collective migration. These elongated streamlines were also less sharply oriented towards the wound and were sometimes dramatically misoriented. In sharp contrast, streamlines from PKN2 KO cells were almost non-apparent in line with the fact that most displacements of PKN2 KO cells were not as collective or persistent as those of parental cells and had a lower amplitude than those of parental cells. This representation can thus become a useful way to capture complex phenotypes.

**Figure 6.**
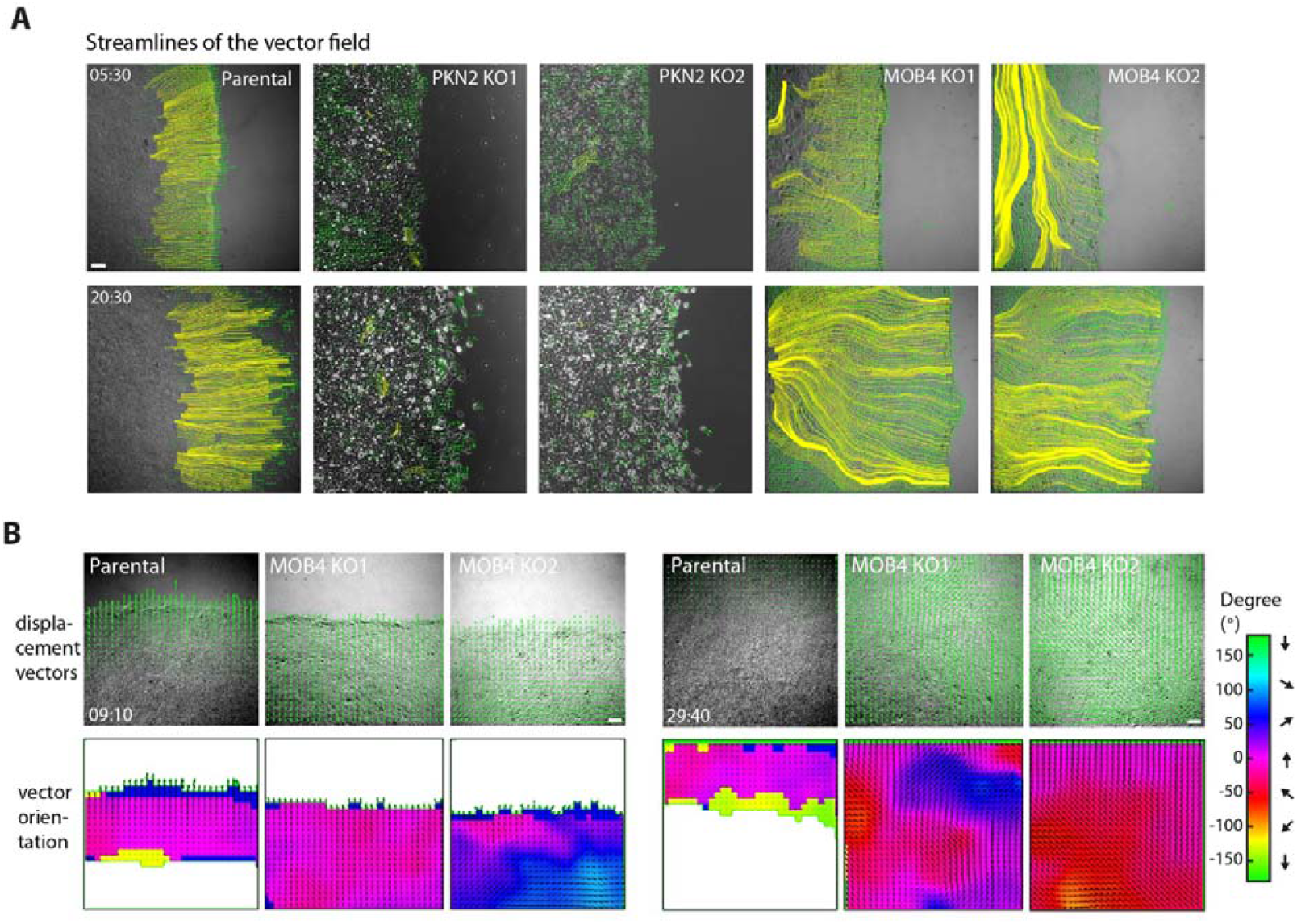
Depletion of MOB4 causes cell streaming in the monolayer. A) Streamlines of the vector field are drawn by concatenating displacement vectors from the beginning of recordings. B) Angular distribution of displacement vectors. Displacement vectors obtained by PIV are superimposed with phase contrast images. Displacement vectors with an amplitude of less than 1 μm/frame are filtered out. Time is in h:min after wounding. Scale bars 100 µm.

Another way to capture the misorientation of collectively migrating MOB4 KO cells is to represent the angular distribution of displacements. Indeed, upon color-coding the angle that displacement vectors make with the overall direction of migration towards the wound, erratic migration directions and unexpected changes of direction of MOB4 KO cells were highlighted (Figure 6B, Figure S9, Video S10). Finally we found that collective migration of MOB4 KO cells was sustained longer than in parental cells: jamming took about 45 h in MOB4 KO clones compared with only 32 h in the control (Figure S9).

Since MOB4 had been described as a non-canonical regulator of the YAP1 pathway ^[58]^, we stained MOB4 KO cells and parental cells with YAP1 antibodies. In a sparse culture, most of the control cells possess nuclear YAP1, whereas YAP1 localization was mostly cytoplasmic in MOB4 KO cells (Figure S10). In contrast, PKN2 KO cells did not show any alteration of YAP1 localization. Upon wounding, front cells of MCF10A cells exhibited YAP1 in the nucleus. This nuclear translocation was particularly clear in the first row of cells next to the wound and declined further back, in the monolayer that remained jammed. In sharp contrast, YAP1 was not translocated in the nucleus in the first rows of cells facing the wound in MOB4 KO cells (Figure 7A). Nuclear translocation of YAP1 is antagonized by its phosphorylation, which is required for its cytosolic retention ^[71,72]^. YAP1 was indeed more phosphorylated on residues S127 and S397 in MOB4 KO cells than in parental MCF10A cells (Figure 7B). We wondered if this defect could account for the migration phenotype of MOB4 KO cells and attempted to rescue KO cells by the expression of a constitutively active form of YAP1 that cannot be inactivated by phosphorylation, YAP1 5SA ^[71]^. We checked that this active YAP1 5SA form had indeed a nuclear localization (Figure S10). Then, we isolated clones of MOB4 KO cells that expressed different levels of active YAP1 (Figure 7C) and submitted these new cell lines to the wound healing assay. Strikingly, the expression of YAP1 5SA abolished the complex MOB4 KO complex phenotype, i.e. increased collective migration with misorientation, in a dose-dependent manner (Figure 7D-F, Figure S11, Video S11). Together, these results show that the restoration of a normal YAP1 function is sufficient to rescue the MOB4 KO phenotype and thus suggest that MOB4 controls collective cell migration through YAP1 activation.

**Figure 7.**
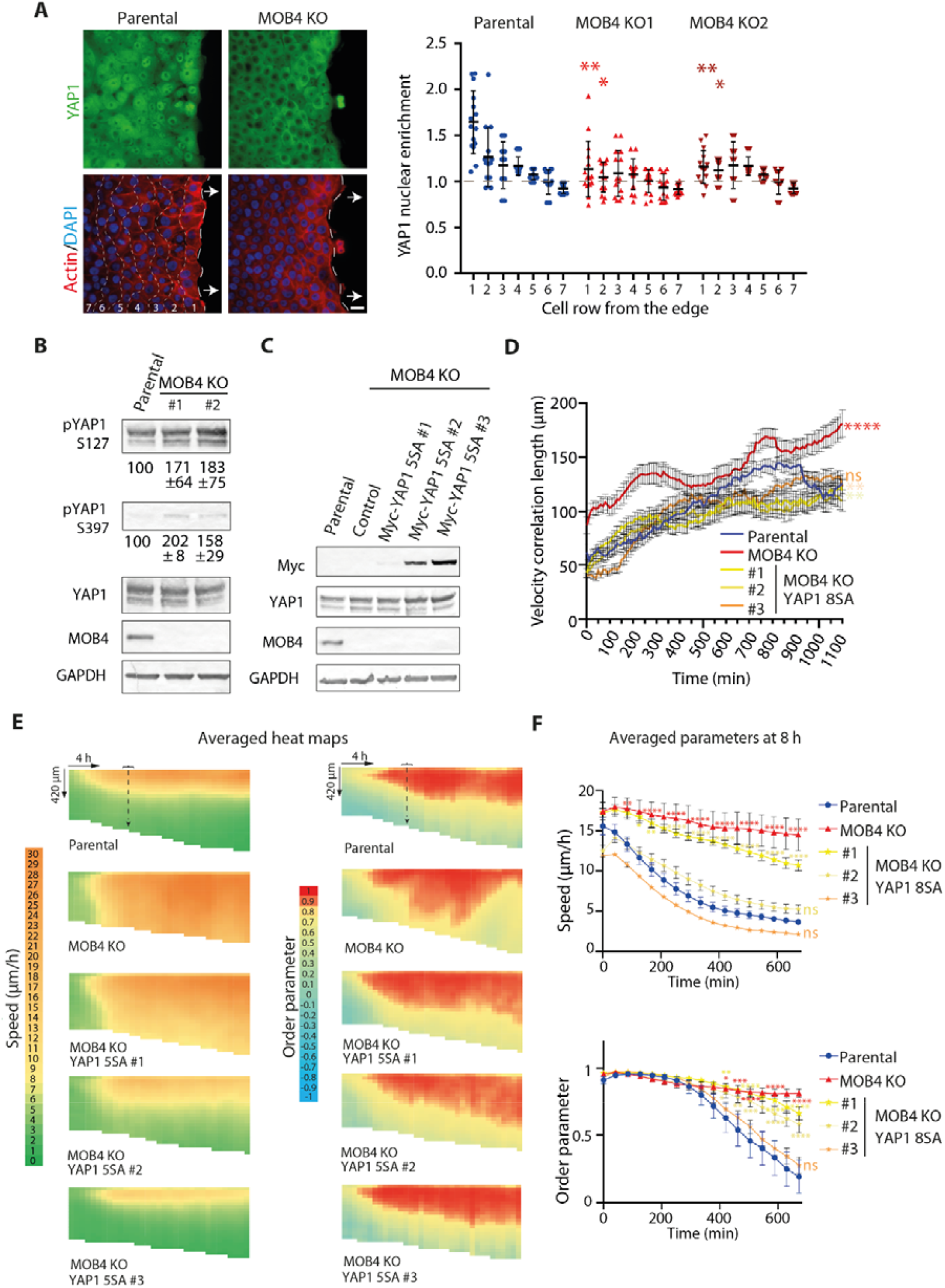
MOB4 regulates collective cell migration through YAP1 activation. A) YAP1 immunofluorescence, phalloidin and DAPI staining in MOB4 KO and parental cells, 8 h after wounding. Wide-field microscopy, scale bar 20 µm. YAP1 nuclear enrichment is assessed in different rows of cells from the wound edge. Five fields of view are analyzed per condition. One-way ANOVA. Three independent experiments gave similar results. A gradient of YAP1 nuclear enrichment from the edge is observed in parental cells, but absent from MOB4 KO cells. B) Western blots of YAP1 and phosphoYAP1 in MOB4 KO and parental cells. The ratio of phosphoYAP1 over YAP1 is indicated; mean ± SD of three independent measurements. C) The constitutively active mutant of YAP1, 5SA, is stably expressed in MOB4 KO cells. Myc Western blots reveal three clones with increasing expression of the YAP1 transgene. D) Velocity correlation length, calculated from 15 movies from three independent experiments, is plotted as a function of time. Mean ± SEM. Areas under the curve are compared by one-way ANOVA. E) Constitutively active YAP1 rescues collective migration of MOB4 KO cells in a dose-dependent manner. Averaged heat maps of 15 movies from three independent experiments are plotted. Each independent experiment gave similar results. F) Speed and the order parameter are averaged at 8 h after wounding from the 15 fields of view of the three biological repeats. Mean ± SEM is plotted, same time points compared with the parental cell line by two-way ANOVA, * p<0.05, ** p<0.01, *** p<0.001, **** p<0.0001. ns, not significant, indicates that no statistical difference was found at any time point with values of parental cells.

## 3. DISCUSSION

We selected five genes encoding proteins associated with cell-cell junctions, among the dependency maps of RAC1-WAVE-Arp2/3. Out of these five candidates, two were indeed involved in the cell coordination of collective migration. Our CRISPR/Cas9-mediated KO approach can be compared to a large siRNA screen similarly performed using wound healing of MCF10A cells ^[73]^. This previous screen identified 66 positive or negative regulators of collective cell migration, out of 1081 genes tested, which included the complete sets of kinases and phosphatases of the human genome and genes identified in another screen for cell adhesion and single cell migration. Our enrichment of candidates appears superior (2/5 >> 66/1081) and therefore allows a careful field analysis of displacement vectors in each KO cell line. Interestingly, the kinase PKN2 was missed in the previous siRNA screen even though other genes of this class, characterized by cell detachment at the front edge of the monolayer, were identified. We report here that cell-cell junctions are destabilized upon PKN2 KO, as measured by the rupture force of E-cadherin coated beads interacting with cells. The dramatic behavior of MOB4 KO cells we report here was to our knowledge never reported before, even if many negative regulators of collective migration, like MOB4, were identified in the published siRNA screen.

We were able to ascribe the complex phenotype of MOB4 KO cells to defective YAP1 activation, since constitutively active YAP1 rescued the MOB4 KO phenotype. Yet all three remaining proteins, DLG5, TP53BP2 and AMOTL2, which did not score positive in our assay, were also YAP1 regulators ^[57,62,74–77]^. MOB4 is a non-canonical regulator of YAP1 activity. Based on published literature, there are several possible ways by which MOB4 can regulate YAP1 activation. MOB4 is a subunit of the striatin-interacting phosphatase and kinase (STRIPAK) complex, a multiprotein complex that contains the PP2A phosphatase ^[60]^ ; in addition to the phosphatase, MOB4 can recruit the MST4 kinase ^[58]^. PP2A phosphatase activity regulates the canonical activity of the essential MST1 kinase, known as Hippo in *Drosophila* ^[78]^. Furthermore, the MOB4-MST4 kinase complex can exchange subunits with the MOB1-MST1 kinase complex and thus alternates between YAP activating and inactivating complexes ^[58]^. In our study, loss of MOB4 leads to an increased phosphorylation of YAP1 and thereby to its inactivation through retention in a cytoplasm. A recently recognized level of YAP1 regulation involves the formation of distinct biomolecular condensates that can coalesce and mix their signaling components, adding further complexity to the signaling pathway ^[79]^. Whatever the exact mechanism involved, the role of MOB4 as a YAP1 activator appears conserved in evolution, since the *C. elegans* MOB4 ortholog controls the life span of the model worm through YAP1 activation ^[80]^.

One of the most remarkable results of our work is that the two hits we identified interact with different domains of the plasma membrane in epithelial cells. PKN2 interacts with lateral edges of collectively migrating cells, whereas MOB4 interact with their front edge. Both PKN2 and MOB4 are cytosolic in the jammed epithelium and relocalize to their respective edge when the monolayer is mechanically unjammed by the wound. Their behavior is thus opposite to the one of NF2, which decorates the whole belt of adherens junctions when cells are jammed and relocalizes to the cytosol upon unjamming ^[31]^. Since both PKN2 and MOB4 interact with the WAVE complex in their respective domain, they can be specific markers of protrusions involved in affixing the membranes of neighboring cells or translocating the cell forward, respectively.

MOB4 inactivation results in a complex phenotype, corresponding to an overall increase of collective migration and the misorientation of this collective migration, a sort of planar cell polarity defect. It is striking that YAP1 activation, and probably the target genes it induces, fully rescue the phenotype. Indeed, since MOB4 is localized in such a way that it can indicate the migration orientation, it is surprising that one of the YAP1 target genes can indicate in the YAP1 rescue experiment where to migrate in the absence of MOB4. In this case, MOB4 and the YAP1 target genes would be redundant in indicating the proper orientation. Another hypothesis might be that MOB4 primarily represses the collective migration within the monolayer through YAP1 and that as a consequence, misorientation of collective migration ensues, because collectively migrating cells are too far to receive the polarity signal from leader cells facing the wound. Understanding how YAP1 target genes contribute to orienting collective cell migration towards the wound is a major challenge ahead.

Our work provides two novel molecular entry points into a complex and finely regulated process. The localization, function and mechanism of PKN2 and MOB4 in collective cell migration were characterized in detail and are summarized in figure 8. Most importantly, these results raise several new questions for future studies. How are different membrane domains specified at the front and on the lateral sides of follower cells, which have neighboring cells all around? How is the polarity signal transmitted from the wound? How is it attenuated from one cell to the next to produce a sharp gradient? Which YAP1 target genes restrict collective migration? And how do they perform this function away from the wound, while being mostly expressed in cells facing the wound?

**Figure 8.**
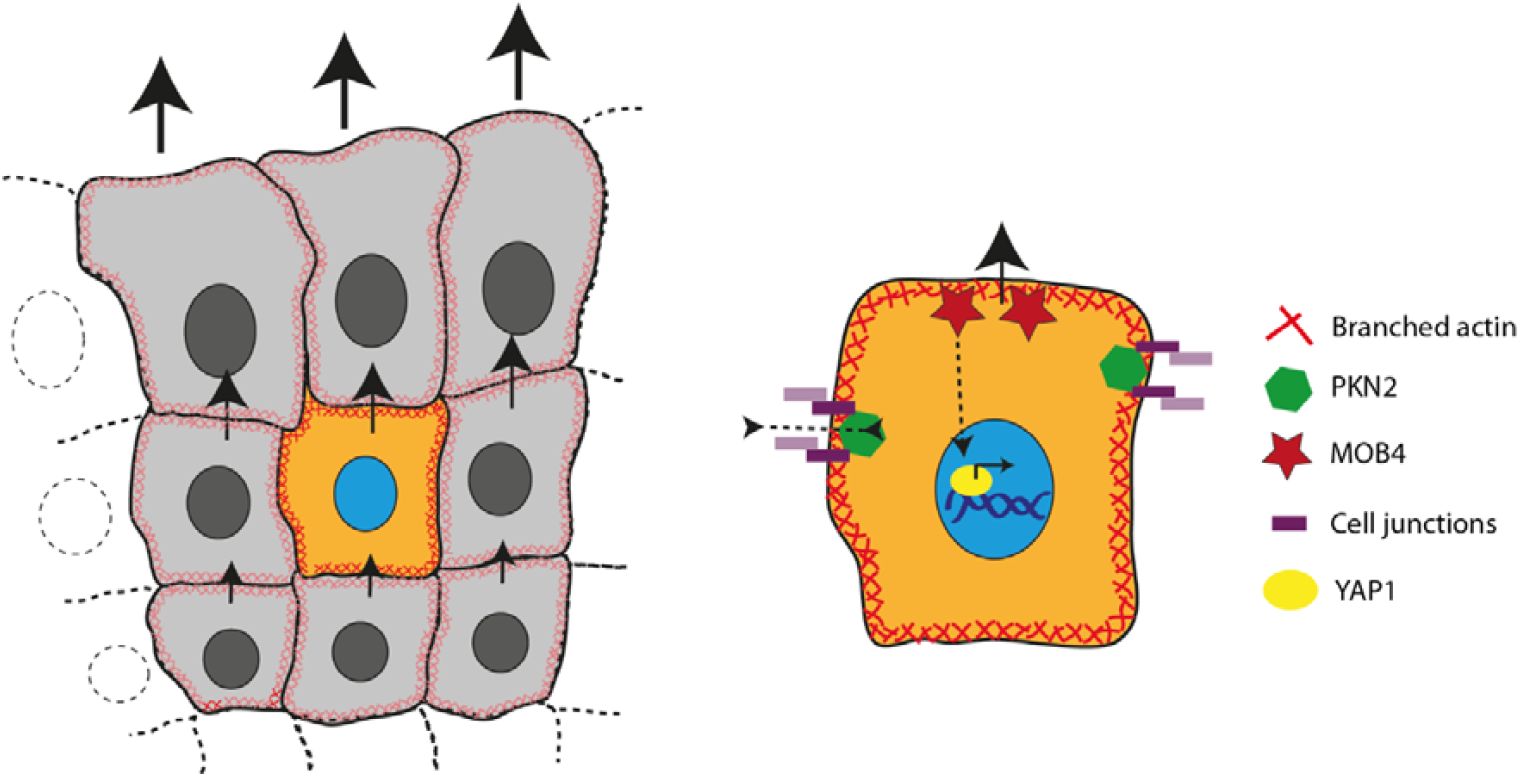
Working model of the roles of PKN2 and MOB4 in the control of collective cell migration. PKN2 localizes at lateral junctions and maintains cohesiveness of the cell monolayer through the maintenance of adherens junctions, whereas MOB4 localizes at the front edge and indicates the cell where to migrate through YAP1 activation. In addition, MOB4 restricts the territory of collectively migrating cells to the first rows of cells facing the wound.

## 4. EXPERIMENTAL SECTION

### Cells

MCF10A cells were a kind gift of Dr. Thierry Dubois, who organizes a collection of mammary epithelial cells of Institut Curie (Paris). They were cultured in DMEM/F12 medium (Gibco) supplemented with 5% horse serum (Sigma), 100 ng/mL cholera toxin (Sigma), 20 ng/mL epidermal growth factor (Sigma), 0.01 mg/mL insulin (Sigma), 500 ng/mL hydrocortisone (Sigma) and 100 U/mL penicillin/streptomycin (Gibco).

### Plasmids and Isolation of Stable Cell Lines

The ORFs of PKN2 (NM_006256) and MOB4 (NM_015387) were amplified from MCF10A cDNA using RNA Plus (Macherey-Nagel) and SuperScript™ III Reverse Transcriptase (ThermoFisher Scientific) Kits. Both cDNAs were cloned into a custom MXS AAVS1L SA2A Puro bGHpA EF1 Flag EGFP Blue SV40pA AAVS1R vector using FseI and AscI restriction sites flanking Blue cassette. The pQCXIH-Myc-YAP-5SA plasmid encoding constitutively active Myc-tagged YAP1 (Uniprot entry P46937-2) comprising the 5 serine to alanine substitutions S61A, S109A, S127A, S164A, and S381A was a gift from Kun-Liang Guan (Addgene plasmid #33093). To obtain stable cell lines, cells were transfected with Lipofectamine 3000 (ThermoFisher Scientific), then diluted and selected with either 1 μg/ml of Puromycin (PKN2 and MOB4) or 15 μg/ml of Hygromycin (YAP1). Individual clones were picked up using cloning rings, expanded and analyzed by Western blot.

### Generation of KO Cell Lines

gRNA sequences were selected in Genome Browser (https://genome-euro.ucsc.edu) based on the best on/off-targeting ratio. Four gRNAs per gene were chosen and their cleavage efficiency was assessed using the T7E1 assay ^[81]^, the most efficient pair at inducing a genomic deletion between the two gRNAs was chosen. Oligonucleotides encoding the corresponding gRNA sequences were then cloned into the pRG2-GG vector at the BsaI restriction site. The gRNA sequences used for each gene were the following: PKN2 gRNA#1 5’-TTGGAGTATCTGGAGTCCTT-3’, PKN2 gRNA#2 5’-GTATTCAAATGGATCTTCAA-3’; TP53BP2 gRNA#1 5’-TCGCCTCTATAAGGAGCTGC-3’, TP53BP2 gRNA#2 5’-CAGCTAGAGATGCTCAAGAA-3’; DLG5 gRNA#1 5’-TTCCACAGGCACTACCGGGA-3’, DLG5 gRNA#2 5’-CAATTACTACCTCTTGTCAA-3’; AMOTL2 gRNA#1, 5’-GGGGGACCGAGATCCCCGTG-3’, AMOTL2 gRNA#2 5’-GGAGCTGCCCACCTATGAGG-3’; MOB4 gRNA#1 5’-AAATGTTTCATTCTGTAGAG-3’, MOB4 gRNA#2 5’-CACATGCTTATTTTCATCAT-3’; gRNA ATP1A1 5’-GTTCCTCTTCTGTAGCAGCT-3’. Efficiency of each cloned gRNA was confirmed by the T7E1 DNA mismatch assay. To amplify the corresponding genomic regions encompassing Cas9-mediated DSBs, the following primer pairs were used: PKN2 for 5’-TCGAACTCAGCTGGAAACAA -3’, PKN2 rev 5’-TTTTTGCATGAATTGGTTGAA -3’; TP53BP2 for 5’-GGAAGAAGGTGAGACCAGACC -3’, TP53BP2 rev 5’-GCACCTGTAACCCCAGCTAC -3’; DLG5 for 5’-CATCTGTGAAAGGGGAATGC -3’, DLG5 rev 5’-GCATTTCTGGACATGAGGTG -3’; AMOTL2 for, 5’-GTACTGAGGTGGGGGTCCTC -3’, AMOTL2 rev 5’-CTGGAACTGGGGGTACAGG -3’; MOB4 for 5’-TCGCTTATTTTCCCTAGGTGTC -3’, MOB4 rev 5’-CCCTTCATGCTTCACTTTCC -3’. To generate KO lines, MCF10A cells were cotransfected using Lipofectamine 3000 (ThermoFischer Scientific) with plasmids encoding Cas9 and encoding the three gRNA sequences, two targeting the specific gene of interest, and the one targeting the *ATP1A1* gene. After 3 days, cells were selected with 0.5 μM ouabain for 5 days. Individual clones were picked after 2 weeks of selection using cloning cylinders. To obtain a control cell line, in addition to parental cells, a stable pool of MCF10A cells resistant to ouabain, with no other genome alteration, was prepared. The genome of potentially KO clones was characterized by PCR using the previously indicated primers. The PCR product was sequenced using Sanger sequencing. If both alleles had the same mutation, the sequence of the PCR product was easily read. In case of overlapping signals in the sequencing chromatogram, the mixed PCR product was cloned into the Zero Blunt vector (ThermoFischer Scientific) and clones corresponding to individual alleles were sequenced.

### Antibodies

The following commercial antibodies were used: DLG5 (ThermoFisher Scientific #A302-302A-M), PKN2 (Proteintech #14608-1-AP), AMOTL2 (Proteintech #23351-1-AP), TP53BP2 (Proteintech #26550-1-AP), MOB4 (Proteintech #15886-1-AP), E-cadherin (clone DECMA-1, Sigma-Aldrich #MABT26), CYFIP2 (Sigma-Aldrich #SAB2701081), NCKAP1 (Bethyl Laboratories, A305-178A), Abi1 (ThermoFischer Scientific #PA5-78705), YAP1 (clone 63.7, SantaCruz Biotechnology #sc-101199), phospho-YAP1 (Ser127) (Proteintech #80694-2-RR, PtgLab), phospho-YAP1 (Ser397) (Proteintech #29018-1-AP), GFP (clones 13.1 and 7.1, Sigma-Aldrich #11814460001), Tubulin (clone DM1A, Sigma-Aldrich #T9026), p150^Glued^/DCTN1 (BD Biosciences #BD610474), GAPDH (ThermoFisher Scientific #AM4300). The following home-made rabbit polyclonal antibodies were used: Myc, WAVE2 and CYFIP1 ^[82]^.

### Western Blots

Cell lysis was performed in 50 mM Hepes, pH7.7, 150 mM NaCl, 1 mM CaCl_2_, 1% NP40, 0.5% Na Deoxycholate, and 0.1% SDS supplemented with protease and phosphatase inhibitor cocktails (Roche). Lysates were centrifuged at 20000 x g at 4°C, supernatants were mixed with LDS loading buffer (ThermoFischer Scientific) and 2.5% of β-ME. SDS-PAGE was performed using NuPAGE 4– 12% Bis-Tris gels (ThermoFischer Scientific), proteins were transferred to a nitrocellulose membrane. Membranes were then blocked in 5% skimmed milk, incubated with primary and secondary antibodies. Secondary antibodies used were conjugated with either alkaline phosphatase (Promega) or with horse radish peroxidase (Sigma-Aldrich). Membranes were developed by using either NBT/BCIP as substrates (Promega) or with SuperSignal™ West Femto Maximum Sensitivity Substrate (ThermoFischer Scientific).

### Immunofluorescence and Image analysis

Cells were seeded on glass coverslips previously coated with 20 µg/mL bovine fibronectin (Sigma) for 1 h at 37°C. Cells were fixed either in sparse culture or jammed or after 8 hours of collective migration depending on the experimental conditions in PBS/3.2% PFA then quenched with 50 Mm NH_4_Cl, permeabilised with 0.5 % Triton X-100, blocked in 2% BSA and incubated with antibodies (1-5 µg/mL for the primary, 5 µg/mL for the secondary) or 1:3000 diluted SiR-actin (Tebu-bio). Secondary goat antibodies conjugated with Alexa Fluor 488, 555 or 647 or DAPI were from Life Technologies/ThermoFischer Scientific. Images were acquired using an inverted Axio Observer microscope (Zeiss) equipped with a 63×/1.4 oil objective, a mercury lamp, and a Hamamatsu camera C10600 Orca-R2 or a SP8ST-WS confocal microscope (Leica) equipped with a HC PL APO 63x/1.40 oil immersion objective, a white light laser, HyD and PMT detectors. For the analysis of orientation of cell-cell junctions, the angle between the direction of the wound edge and the junction stained with MOB4 or PKN2 was measured. All junctions were assessed in several fields of view. For the E-cadherin intensity profiles, multiple lines were drawn orthogonally to the junction identified by an actin staining and these lines were averaged per junction. For nuclear enrichment of YAP1, cells were thresholded in the DAPI and actin channel so as to saturate nucleus and cytosolic staining. YAP1 intensity was then measured in the regions of interest corresponding either to DAPI (i.e nuclear intensity) or the difference between actin and DAPI (i.e. cytoplasmic intensity). The ratio of both intensities gave the nuclear enrichment.

### Immunoprecipitation

Cells expressing GFP fusion proteins were lysed with XB-NP40 buffer (50 mM HEPES, 50 mM KCl, 1% NP-40, 10 mM EDTA, pH7.7) supplemented with protease inhibitors during 30 min on ice. Clarified cell extracts were incubated with GFP-trap beads (Chromotek) for 1 h at 4 °C. The GFP-trap beads were washed with XB-NP40 buffer. Beads were subjected to SDS–PAGE and Western blot.

### Wound Healing

For wound healing experiments, 80 000 cells were seeded in a Ibidi insert (Ibidi, 80209) and allowed to adhere for a 24 h. Wound was created by removing the insert. Cells were washed with fresh medium and imaged using videomicroscopy. Time-lapse multi-field experiments were performed in phase contrast or in GFP/mCherry fluorescent channels on automated inverted microscopes (Olympus IX71 or Zeiss AxioObserver Zeiss) both equipped with thermal and CO_2_ regulation. Acquisitions were performed using a charge-coupled device (CCD) camera (Retiga 4000⍰R, QImaging or Hamamatsu camera C10600 Orca-R2) and Metamorph (Universal Imaging) or Micro Manager (ImageJ) software to control them. Images were taken every 10 min.

Phase contrast movies were analyzed using particle image velocimetry (PIV) performed in the PIVlab software package ^[83]^ for Matlab (MathWorks). To obtain the field of displacement vectors, the window size was set to 32 pixels⍰=⍰23.75⍰µm with a 0.75 overlap. Spurious vectors were filtered out by their amplitude and replaced by interpolated velocity from neighboring vectors. A time sliding window averaging velocity fields over 40 minutes (4 frames) was used. Subsequent analysis was performed as previously described ^[67]^ using the published code to extract the order parameter and cell speed. Robustness of the analysis was checked by using a time of 30 min between successive images. Velocity correlation length was extracted from velocity fields as previously described ^[22]^. To draw streamlines and represent angular orientations, displacement vectors with an amplitude less than 1 μm/frame were filtered out.

### Single Cell Migration and Aggregation Assays

For both assays, cells were plated in 8-well Slides (Ibidi, 80806) coated with Fibronectin and allowed them to adhere during 24 hours. Cells were then imaged during 48 h in phase contrast on Zeiss AxioObserver, images were taken every 10 minutes. Cells were tracked manually in FIJI and parameters of single cell migration were extracted from cell trajectories as described previously (Gorelik and Gautreau, 2014). To assess cell aggregation, number of cells and cell islets were quantified manually at different time points.

### Force Measurement Using E-cadherin Coated Beads

We employed a procedure that we had previously set up ^[70]^. In brief, experiments were conducted using an inverted epifluorescence microscope (Olympus Live EZ) equipped with a 40× objective (LUCPLFLN; NA/0.6) and an Andor Neo 5.5 CMOS (2560 x 2160px). The cells were seeded in a chamber composed of a U-shaped PDMS structure with two glass coverslips—one on top and one at the bottom. The bottom coverslip was coated with fibronectin to facilitate cell attachment. Amine Dynabeads M270 (ThermoFischer Scientific #14307D) were conjugated with purified E-cadherin extracellular domain (SinoBiological #10204-H02H). The control corresponded to the absence of E-cadherin extracellular domain. The efficiency of cadherin coating was assessed in a bead aggregation assay. To this end, equal number of E-cadherin-coated Amine Dynabeads M270 (2.7⍰μm diameter) were resuspended and incubated at 37°C for 5 min in 1⍰mL of 20⍰mM Tris-HCl, pH 7.4, supplemented either with 1⍰mM CaCl_2_ or 1⍰mM CaCl_2_ + 4⍰mM EDTA. Microbeads were then gently pelleted by centrifugation at 500⍰rpm for 30 s and incubated for an additional 5 min at 37°C to promote bead–bead interactions and were subsequently resuspended by gently inverting the tube five times. Then images were acquired in brightfield mode and aggregated beads (defined as clusters of two or more beads) were counted using ImageJ. The aggregation ratio was calculated as:

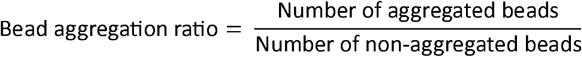

E-cadherin coating was efficient, since bead aggregation ratio was 0.35 ± 0.14 in Ctrl, 1⍰mM CaCl_2_ increased it to 0.67 ± 0.09 (p<0.001) and EDTA addition reverted it to 0.28 ± 0.08 (non-significant in paired t-test).

Elastic micropipettes were created by pulling them (Sutter P-97 micropipette puller) and cutting them with a homemade microforge to achieve an inner dimension of 2-3 μm. The micropipette was hold by a micropipette holder attached to a micromanipulator (Sutter MP-285), and the system was connected to a hydraulic setup to control bead capture through aspiration. The micropipette was maneuvered to contact the cell edge and gently pushed towards the cell with a displacement of 1-2 μm. Subsequently, the micropipette was gradually withdrawn from the cell by moving the chamber’s stage in 3 μm increments at an average speed of 5 μm/s until detachment occurred. The adhesive interaction between the bead and the cell was specifically ruptured. The bending deflection of the micropipette (x) was recorded, and the micropipette stiffness (k) was determined following calibration with a standard micropipette. The rupture force was calculated using the formula F = kx. Mean force values after 5 min of contact were derived from the results of more than 10 cells in each group, obtained from at least three independent experiments.

### Statistics

Statistical analyses were carried out with GraphPad Prism software (v7.00) and Microsoft Excel 2016. Shapiro-Wilk normality test was applied to examine whether data fit a normal distribution. If data satisfied the normality criterion, ANOVA was followed by a post-hoc Tukey’s multiple comparison test. If not, a non-parametric Kruskal-Wallis test was followed by a post-hoc Dunn’s multiple comparison test. Pairwise comparisons were assessed by either the t-test if data followed a normal distribution or by the Mann-Whitney test otherwise. Four levels of significance were distinguished: * p<0.05, ** p<0.01, *** p<0.001, **** p<0.0001.

## Author Contributions

A.I.F. performed most experiments and data analysis in the two research groups of P.S. and A.M.G. Y.L. performed force measurements using E-cadherin-coated beads. D.Y.G. adapted the ouabain selection procedure to the double cut approach for fast selection of KO cell lines. AIF and DYG isolated and characterized KO cell lines. H.Y.C. helped with PIV and analysis of wound healing; J.J. helped with data analysis; J.Y., P.S. and A.M.G. supervised the study in their respective research group. AIF and AMG jointly wrote the manuscript.

## Conflict of interest

The authors declare no conflict of interest

## Supporting Information

Supplementary information is available.

## Data Availability

The data that support the findings of this study are available in the supplementary information.

## Acknowledgements

This work was supported by grants from Agence Nationale de la Recherche (ANR-20-CE13-0016 to PS and AMG, ANR-22-CE13-0041 and ANR-24-CE44-4957 to AMG), Fondation ARC pour la Recherche sur le Cancer (ARC PJA 2021 060003815 to AMG) and Institut National du Cancer (INCA_16712 to AMG). This work benefited from the support of the Personalized reconstitution of the tumoral process program led by l’X, Ecole polytechnique and the Fondation de l’Ecole polytechnique, sponsored by Servier. We thank the Polytechnique Bioimaging Facility for confocal microscopy partly supported by Agence Nationale de la Recherche (ANR-11-EQPX-0029 Morphoscope2, ANR-10-INBS-04 France BioImaging). Work in Institut Curie received support from the Labex Cell(n)Scales (grants ANR-11-LABX-0038, ANR-10-IDEX-0001-02), and received support from “Institut Pierre-Gilles de Gennes” laboratoire d’excellence, “Investissements d’avenir” program ANR-10-IDEX-0001-02 PSL and ANR-10-LABX-31, from Canceropôle Ile-de-France and French National Cancer Institute. We thank the Cell and Tissue Imaging core facility (PICT IBiSA), Institut Curie, member of the French National Research Infrastructure France-BioImaging (ANR10-INBS-04) and the Molecular Biology and Cellular Biology platform of CNRS UMR168. This study was supported by the Single-cell mechanical study was supported by National Research Foundation (NRF) Singapore, Mechanobiology Institute under its MID-SIZED GRANT (MSG) (NRF-MSG-2023-0001 to J.Y.). We thank Prof. Kun-Liang Guan (UCSD, CA) for the kind gift of the YAP1 plasmid and Dr. Anna Polesskaya for critical reading of the manuscript.

## Notes

### Competing Interest Statement

The authors have declared no competing interest.

### Summary of Updates

-Qualification of the MOB4 KO phenotype: streamlines in Fig.6A introduced to avoid using swirling -Inclusion of a new migration parameter: velocity correlation length in Fig.2D, 5B, 7D -Confocal microscopy to localize PKN2 and MOB4 at cell-cell junctions (Fig.3 and 4) -Important control: no effect of the PKN2 and MOB4 KO on the actin cytoskeleton at cell-cell junctions (Fig.4C) -Inclusion of averaged and single fields of view heat maps for all collective migration results (Fig. S3, S12, S14) -Assays of single cell migration (Fig.S5)

