## Supplemental Information for "Identification of PKN2 and MOB4 as Coordinators of Collective Cell Migration"

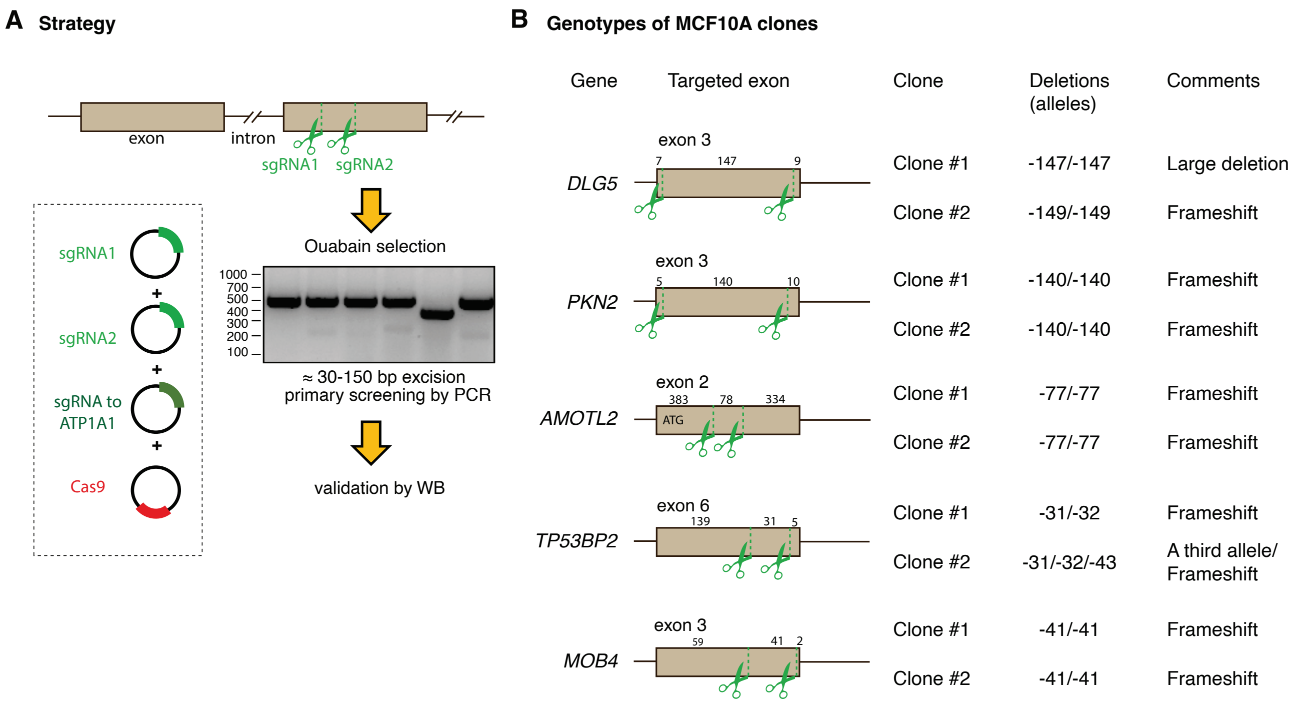


**Figure S1**. **Characterization of MCF10A KO cell lines.** A) Strategy used to generate KOs using ouabain selection. B) Genotypes of KO cell lines. The exact positions of Cas9-induced cuts in each targeted exon are indicated. The majority of clones correspond to the religation after the double cut with no further modification. Some alleles, however, were slightly different, with a couple of extra-nucleotides that were added or removed. The two alleles of a clone most often exhibit the same deletion. Independently isolated clones frequently exhibited the same deletion.

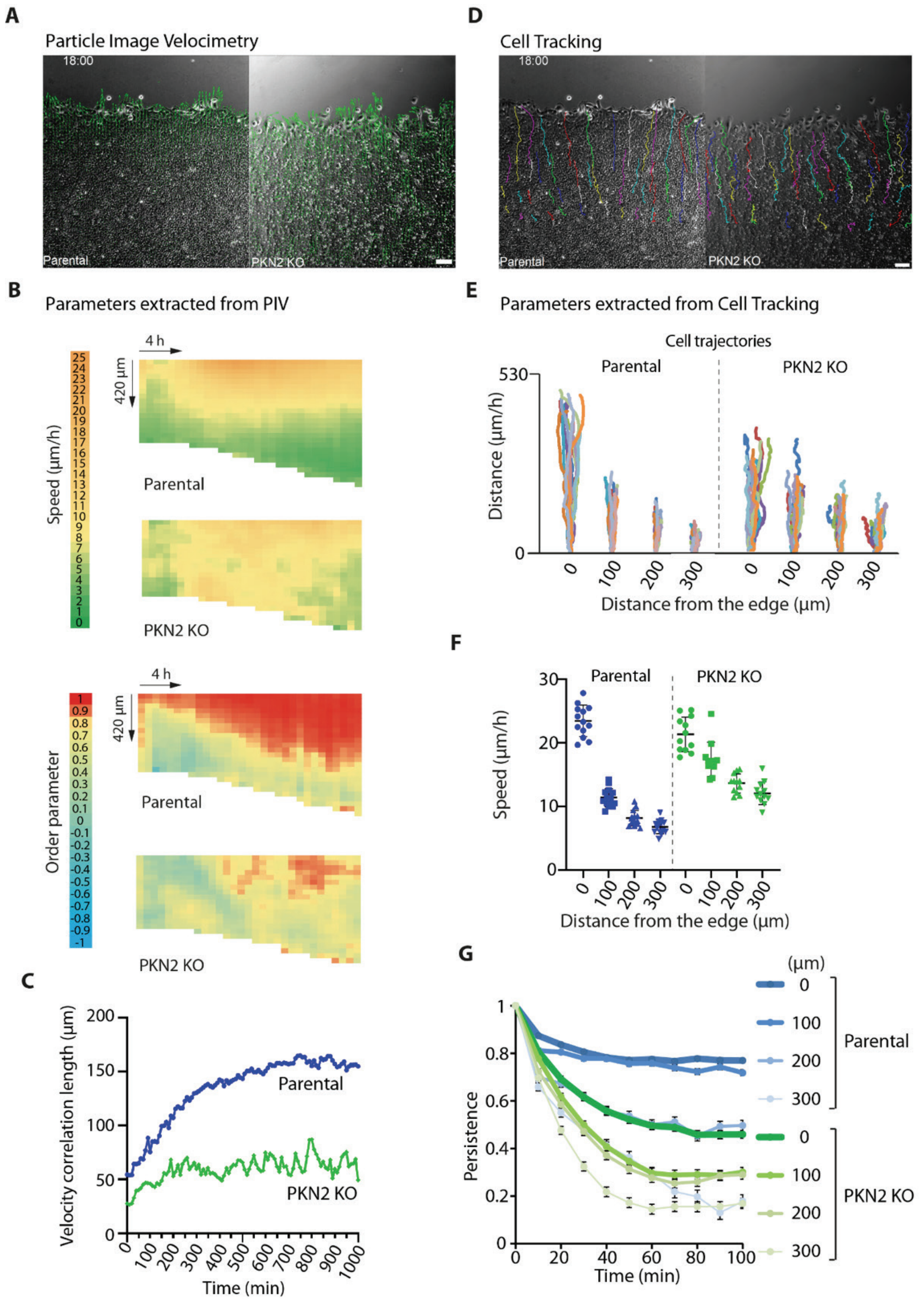


**Figure S2. Collective migration of PKN2 KO and parental cells analyzed using Particle Image Velocimetry (PIV) or Cell Tracking.** A-C) Extraction of migration parameters from PIV analysis of single fields of view. A) Examples of vector fields superimposed with phase contrast images at 18 h after wounding. B) Speed and order parameter plotted as heat maps across time and space. C) Velocity correlation length plotted as a function of time. D-G) Extraction of migration parameters from cell tracking. Migration is analyzed from the trajectories of 13 cells located at four different distances from the wound edge. D) Cell trajectories superimposed with phase contrast images at 18 h after wounding. E) Length of cell trajectories. F) Cell speed, mean ± SEM. G) Directional persistence, corresponding to the autocorrelation of displacement orientation (cosine of the angle between two displacement vectors) over various time intervals, at four different distances from the wound edge. Mean ± SEM.

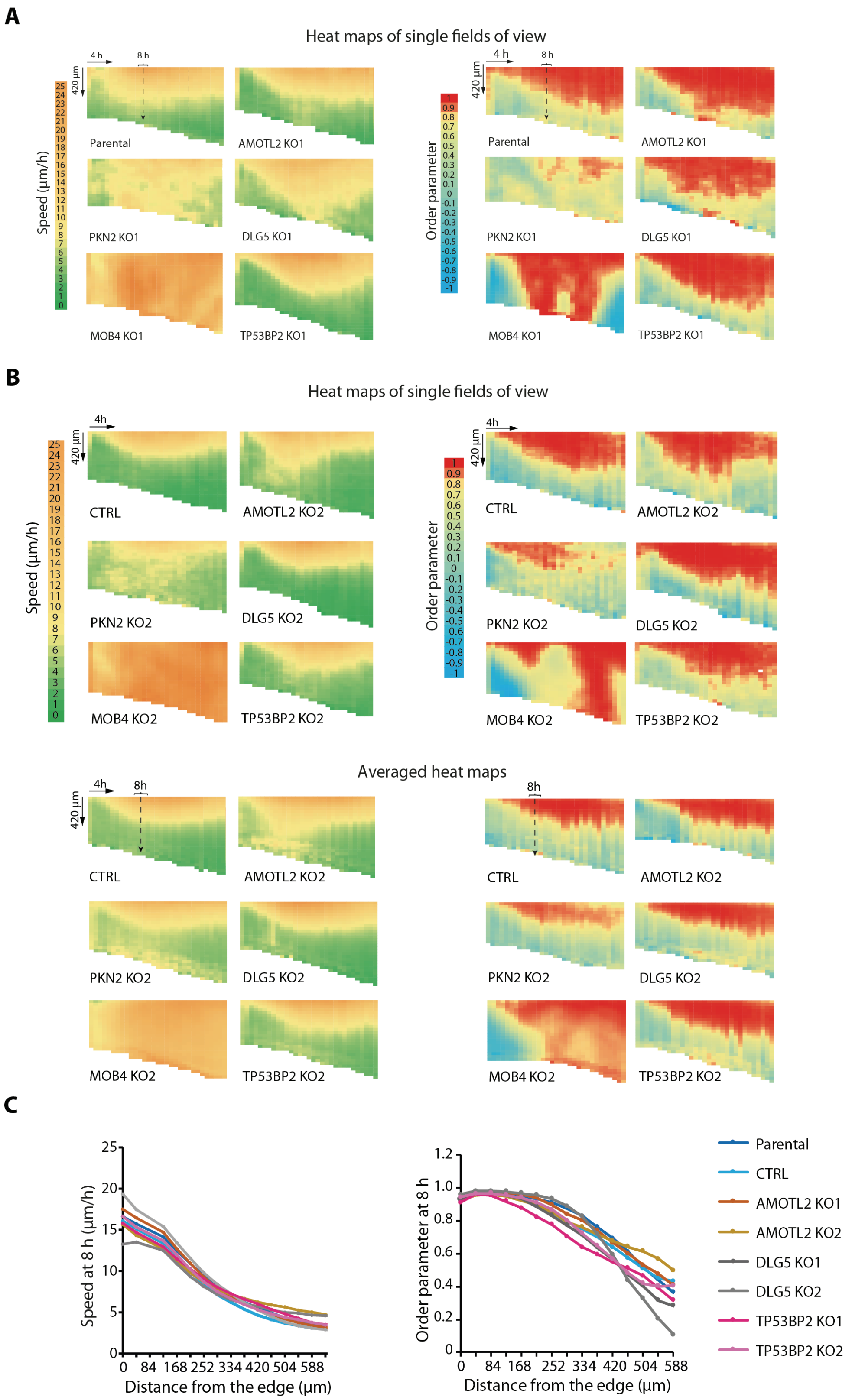


**Figure S3. Further characterization of collective cell migration.** A) Heat maps of speed and the order parameter from single fields of view. B) Heat maps of speed and the order parameter from single and averaged fields of view for a second clone of each genotype. C) Speed and order parameter at 8 h after wounding of the three non-selected KOs. The average is from the eight fields of view of two independent experiments.

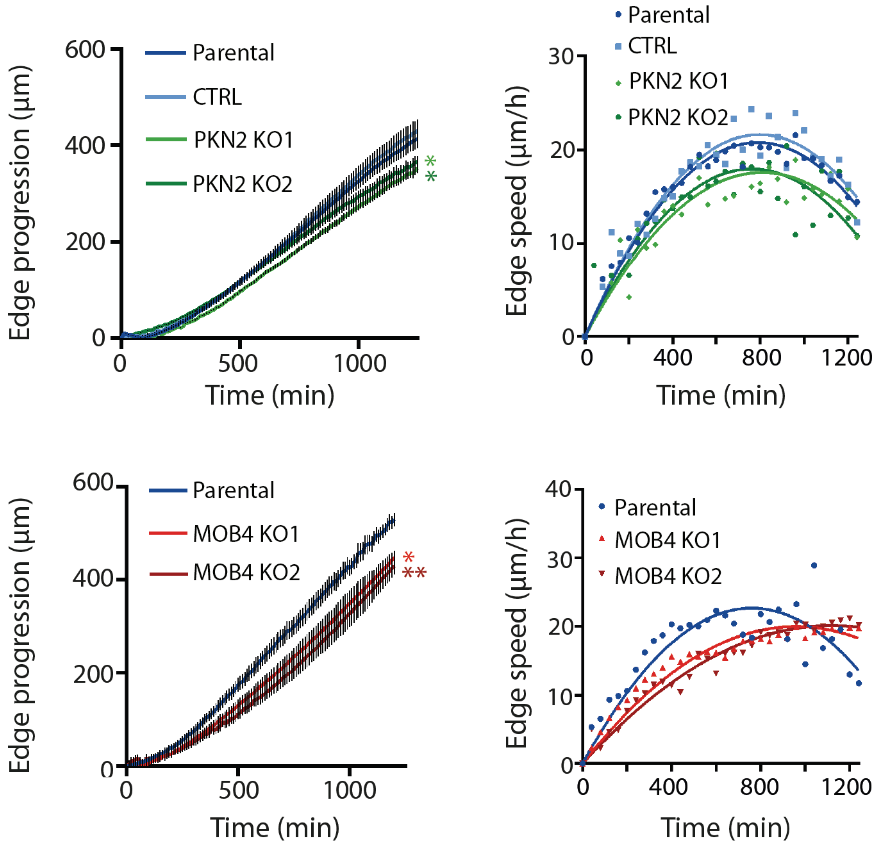


**Figure S4. Wound healing by PKN2 and MOB4 KO cell lines**. Edge progression and speed are plotted as a function of time. Average of 12 fields of view for three independent experiments. * $P<0.05$, ** $P<0.01$.

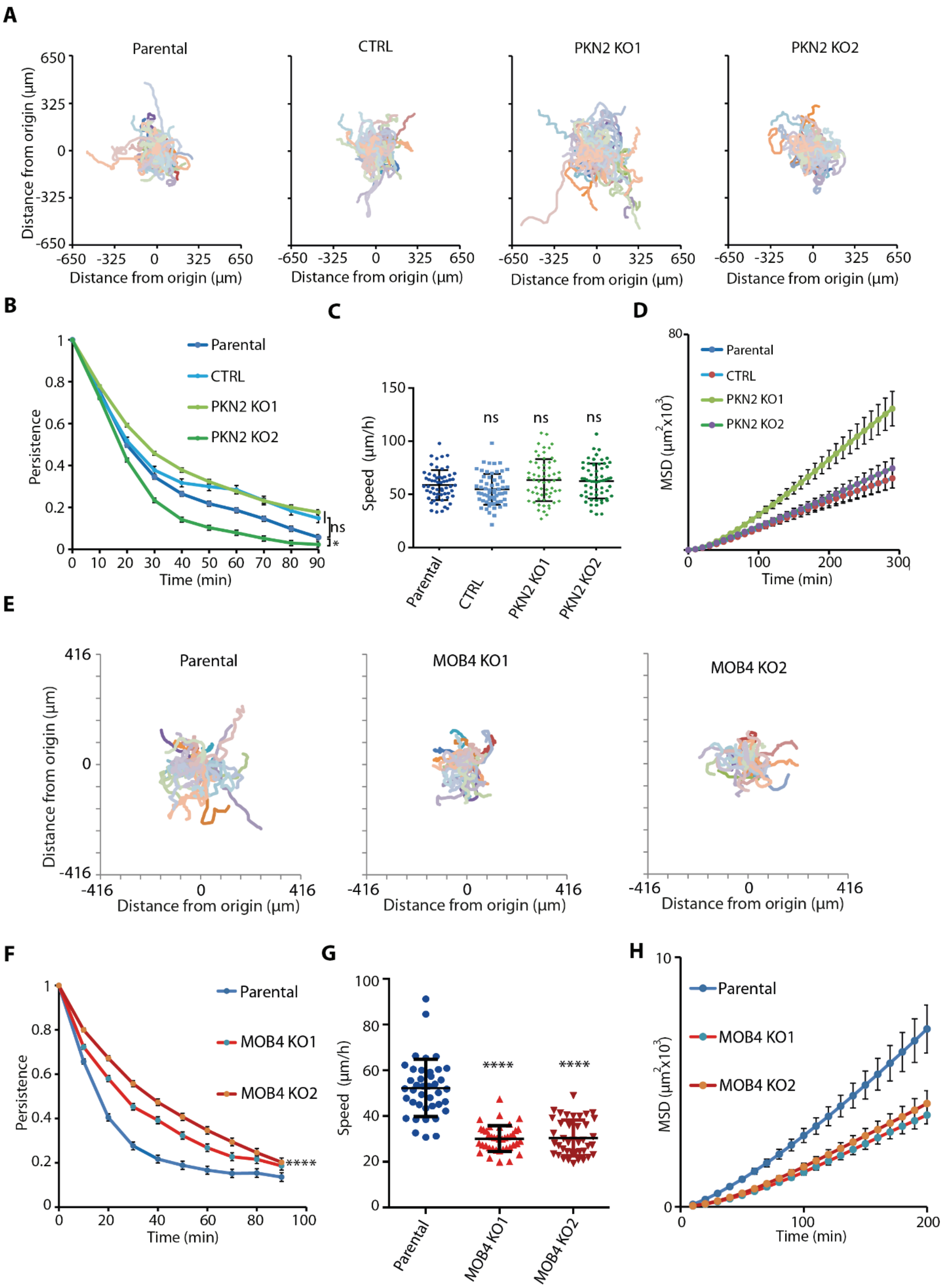


**Figure S5**. **Single cell migration.** A-D) Analysis of PKN2 KO clones and controls. 60 cells from two independent experiments per condition are analyzed. A) Cell trajectories. B) Cell persistence, mean ± SEM. C) Speed, mean ± SEM. D) Mean Square Displacement (MSD), mean ± SEM. E-F) Analysis of MOB4 KO clones and parental cell line. 60 cells from two independent experiments per condition are analyzed. E) Cell trajectories. F) Directional persistence, mean ± SEM. G) Speed, mean ± SEM. H) MSD, mean ± SEM. * $p<0.05$, **** $p<0.0001$, ns not significant.

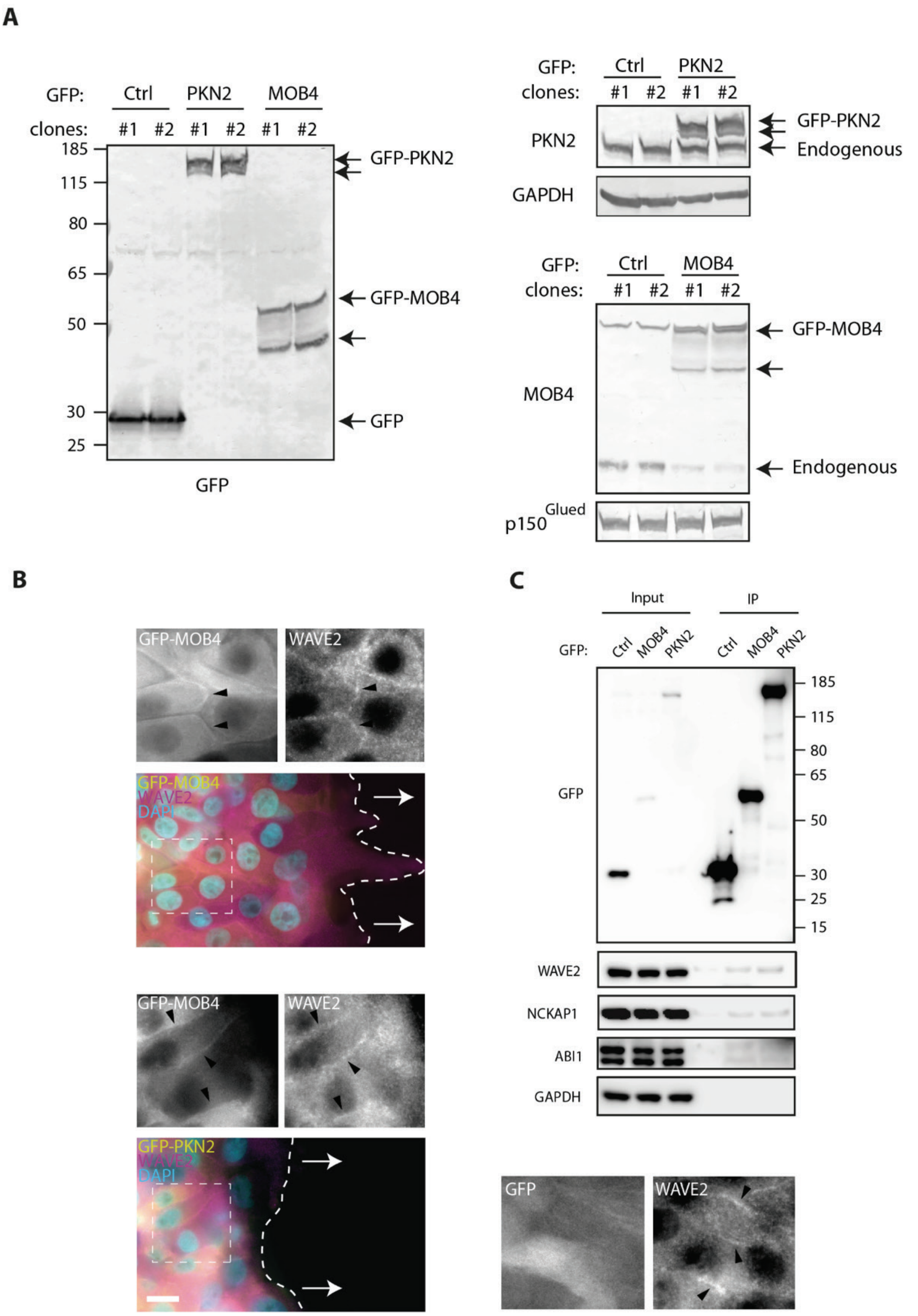


**Figure S6. MOB4 and PKN2 associate with the WAVE complex at cell-cell junctions** A) Characterization of stable MCF10A cell lines expressing GFP-tagged MOB4 and PKN2. Western blots of two clones of the indicated cell lines using MOB4, PKN2 and GFP antibodies. GAPDH and $p150^{Glued}$ are loading controls. Note that tagged MOB4, but not tagged PKN2, replace its endogenous counterpart. B) Both PKN2 and MOB4 colocalize with WAVE2 during collective cell migration, 8 h after wounding. Still images extracted from wide-field videomicroscopy, scale bar 20 µm. C) Immunoprecipitation of GFP-tagged MOB4, PKN2, or GFP as a control. The WAVE complex subunits, WAVE2, NCKAP1 and ABI1, co-precipitate with both MOB4 and PKN2. One experiment representative of three is shown.

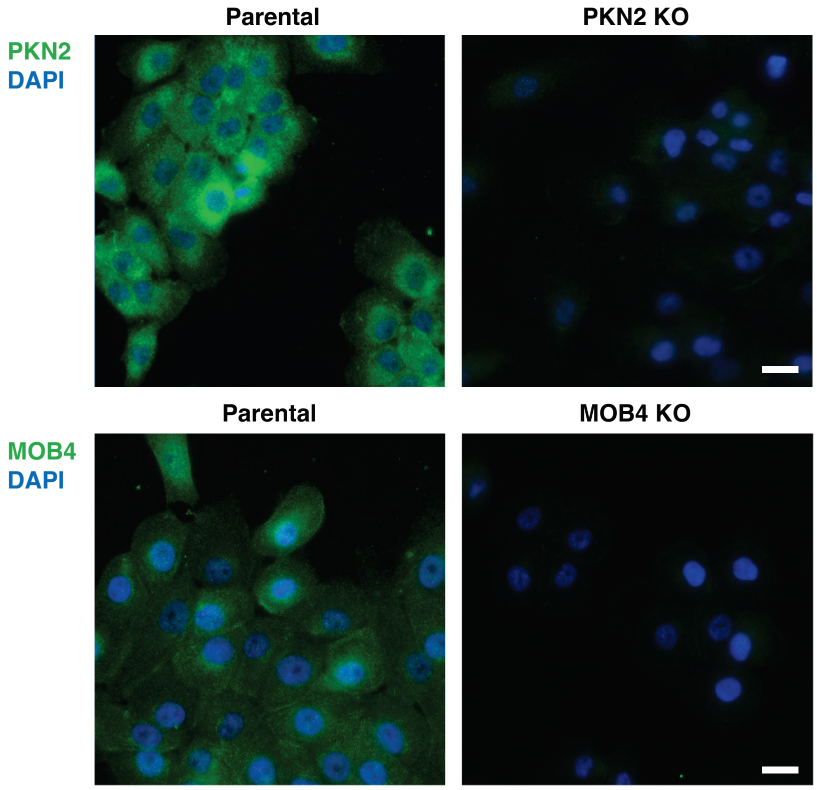


**Figure S7. Specificity of PKN2 and MOB4 antibodies.** Parental or KO cells are stained by immunofluorescence using corresponding antibodies and imaged by wide-field microscopy using the same exposure time. Scale bar 20 µm.

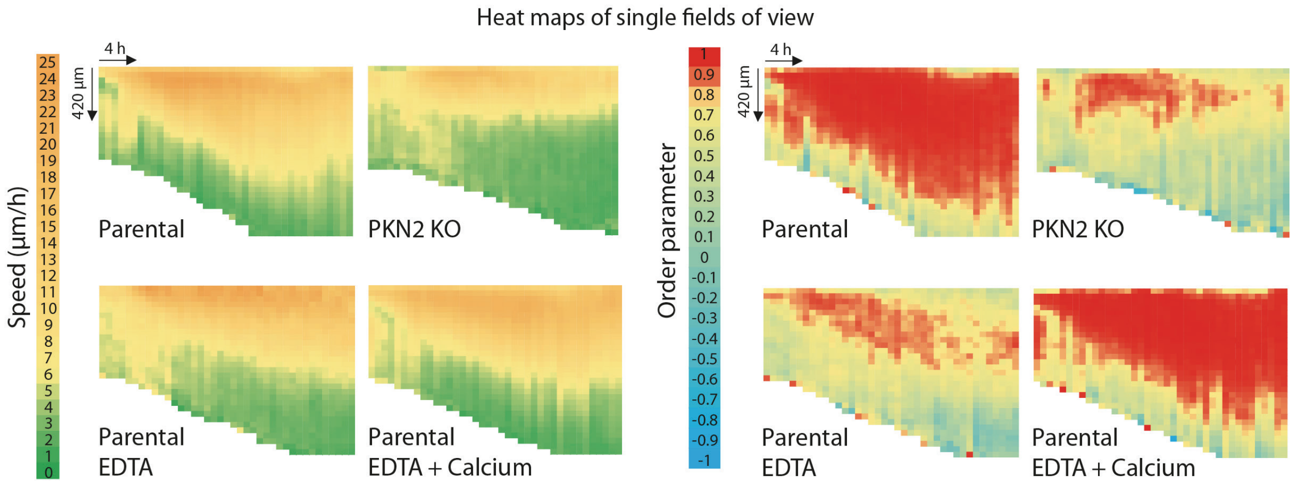


**Figure S8. Speed and order parameter of PKN2 KO and EDTA treated cells.** Heat maps are extracted from single fields of view.

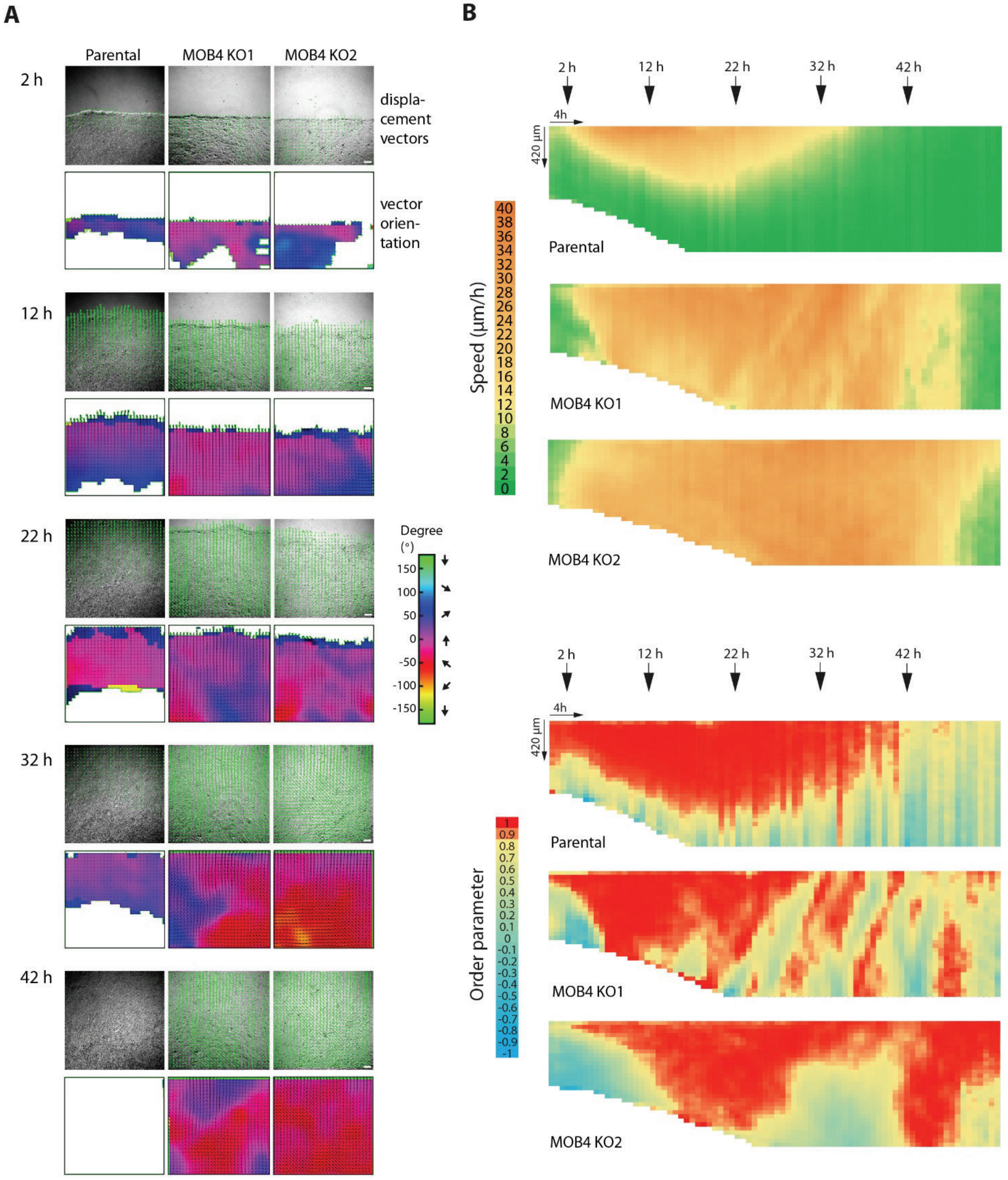


**Figure S9. MOB4 KO cells collectively migrate for a longer time than parental cells, but not always towards the wound.** A) Angular distribution of displacement vectors. Displacement vectors obtained by PIV are superimposed with phase contrast images. Time in h after wounding. B) Speed and the order parameter of MOB4 KO cells over an extended time course. Heat maps are generated from a single field of view. Similar results were obtained in two independent experiments with four fields of view. MOB4 KO cells take longer to jam than parental cells.

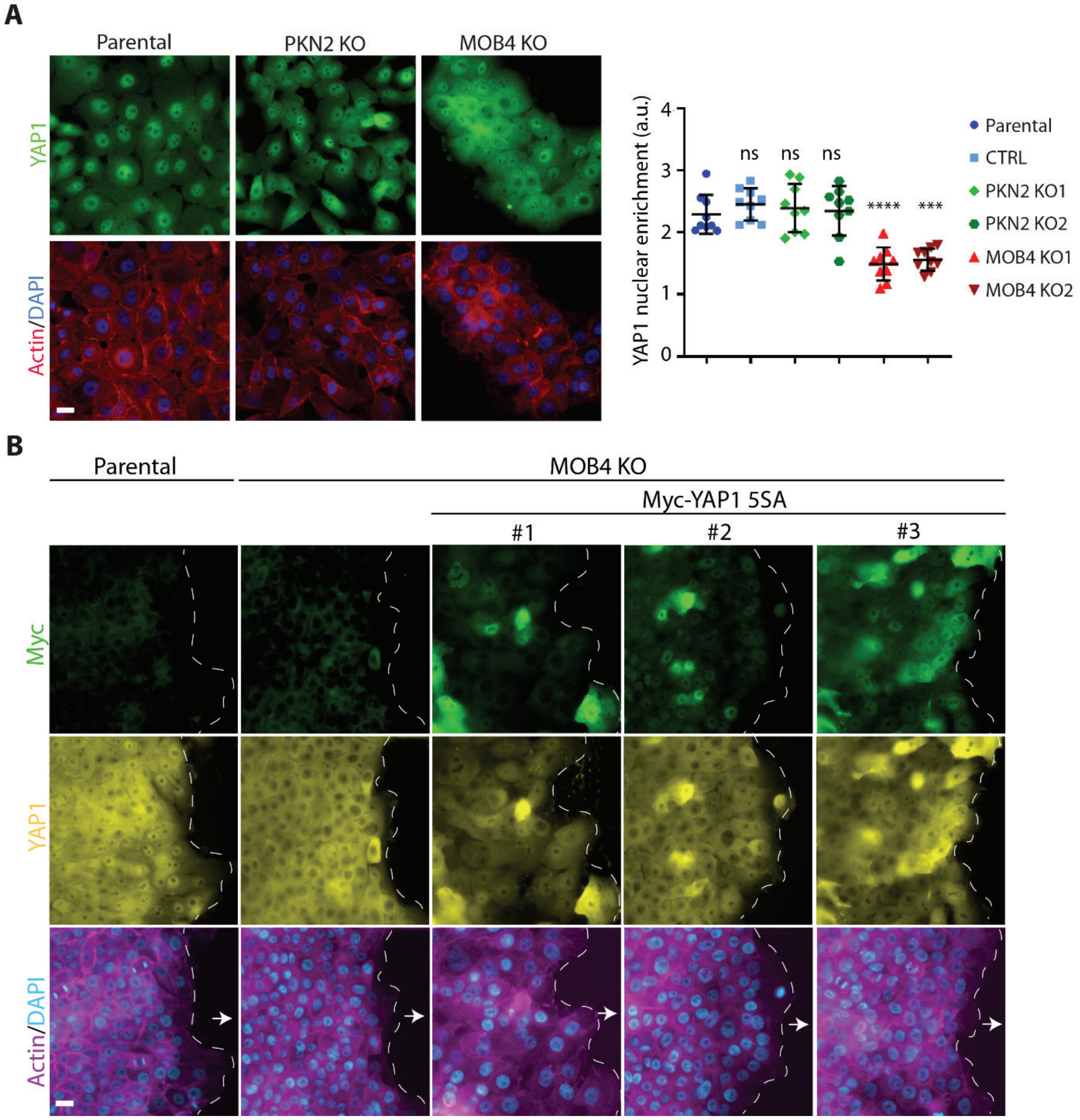


**Figure S10**. **YAP1 in MOB4 KO cells.** A) Immunofluorescence of YAP1. PKN2 KO, MOB4 KO and parental cells are stained with YAP1 antibodies, phalloidin and DAPI. Wide-field microscopy, scale bar 20 µm. YAP1 nuclear enrichment is calculated in nine fields of view per condition from three independent experiments. Mean ± SD. One-way ANOVA, *** p<0,001, **** p<0.0001, ns non-significant. B) The constitutively active YAP1 5SA localizes to the nucleus. MOB4 KO or parental cells transfected with the construct as indicated are wounded. Cells are fixed 8 h after wounding and stained with antibodies targeting Myc, antibodies targeting YAP1, phalloidin and DAPI. Wide-field microscopy, scale bar: 20 µm.

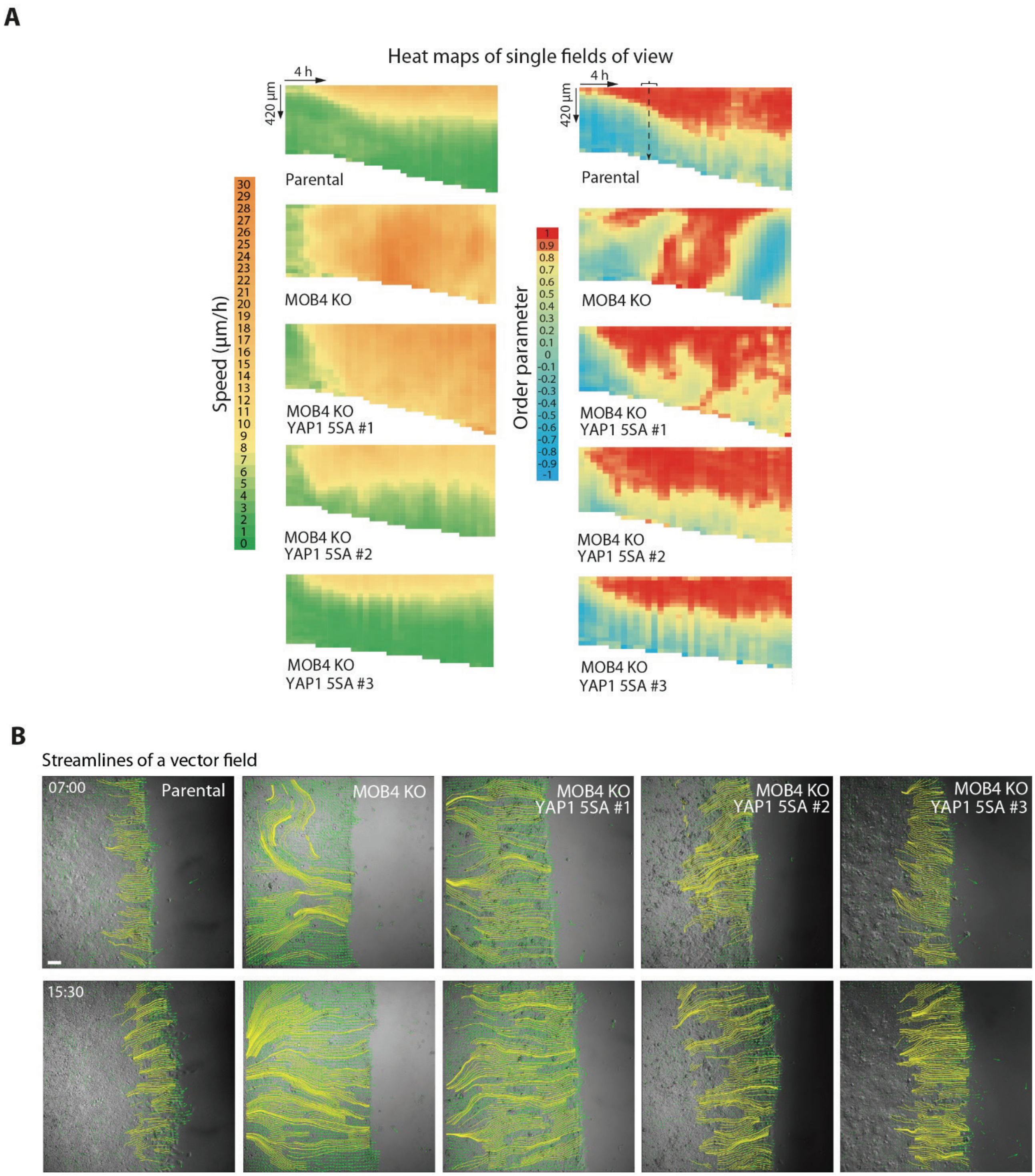


**Figure S11. Constitutively active YAP1 rescues the phenotype of MOB4 KO.** A) Individual representative heat maps of collective cell migration. B) Streamlines of displacement vectors. Time in h:min after wounding. Scale bar 100 µm.

**Movie Legends**

**Video S1. Wound healing of PKN2 KO cells with PIV displacement vectors.** A monolayer of PKN2 KO or parental cells is wounded at time 0. Healing is recorded by videomicroscopy using phase contrast. Displacement vectors obtained by PIV are superimposed to phase contrast images. Timer in h:min. Scale bar 100 µm.

**Video S2. Wound healing of PKN2 KO cells with single cell tracking.** Tracking of single PKN2 KO and parental cells within the wounded monolayer. Trajectories of front cells at the leading edge or distant from the edge by 100, 200 and 300 µm are superimposed with phase contrast images. Timer in h:min. Scale bar 100 µm.

**Video S3. Wound healing of MOB4 KO cells.** A monolayer of MOB4 KO or parental cells is wounded at time 0. Healing is recorded by videomicroscopy using phase contrast. Displacement vectors obtained by PIV are superimposed to phase contrast images. Timer in h:min. Scale bar 100 µm.

**Video S4. Individual examples of wound healing by MOB4 KO cells.** Three fields of view of the same MOB4 KO clone or parental cells are wounded at time 0. Healing is recorded by videomicroscopy using phase contrast. Displacement vectors obtained by PIV are superimposed to phase contrast images. Timer in h:min. Scale bar 100 µm.

**Video S5. Single cell migration of PKN2 KO and parental cells.** Single cells are tracked and trajectories are superimposed with phase contrast images. Timer in h:min. Scale bar 100 µm.

**Video S6. Single cell migration of MOB4 KO and parental cells.** Single cells are tracked and trajectories are superimposed with phase contrast images. Timer in h:min. Scale bar 100 µm.

**Video S7. GFP-PKN2 and GFP-MOB4 relocalizes to distinct domains of the plasma membrane upon wound healing.** A monolayer of cells stably expressing GFP, GFP-PKN2 or GFP-MOB4, is wounded at time 0. Healing is recorded by wide-field videomicroscopy using GFP fluorescence. Arrowheads indicate membrane domains decorated by PKN2 or MOB4. Timer in h:min. Scale bar 20 µm.

**Video S8. Wound healing by a mixed population of non-fluorescent parental cells and GFP-MOB4 cells.** Cells are counter-stained using the vital dye SiR-Actin. GFP-MOB4 decorates the front, but not the back edge of migrating cells (arrowheads). Wide-field videomicroscopy. Timer in h:min. Scale bar 20 µm.

**Video S9. EDTA-mediated destabilization of junctions in parental cells phenocopies PKN2 KO cells.** Wound healing of PKN2 KO cells or parental cells treated with 1 mM EDTA to destabilize cell-cell junctions or 1 mM EDTA with 1 mM $CaCl_2$ to restore E-cadherin homotypic binding. Timer in h:min and scale bar 100 µm. Note the cells that dissociate from the edge of PKN2 KO monolayer or parental cell monolayer treated with EDTA.

**Video S10. Misorientation of MOB4 KO cells.** Monolayers of the two MOB4 KO clones or parental cells are wounded at time 0. Healing is recorded by videomicroscopy using phase contrast. Displacement vectors obtained by PIV are superimposed to phase contrast images. Their orientation is color-coded according to the scale displayed. Timer in h:min. Scale bar 100 µm.

**Video S11. Active YAP1 rescues MOB4 KO cells.** A cell monolayer of the indicated genotype is wounded at time 0. Healing is recorded by videomicroscopy using phase contrast. Displacement vectors obtained by PIV are superimposed to phase contrast images. Timer in h:min. Scale bar 100 µm.
